# Can Large Language Models “Read” Biological Sequences? A Systematic Evaluation of In-Context Learning for Antibody Characterization

**DOI:** 10.1101/2025.02.11.637772

**Authors:** Sin-Hang Fung, Zhenghao Zhang, Ran Wang, Chen Miao, Brian Shing-Hei Wong, Kelly Yichen Li, Chenyang Hong, Jingying Zhou, Kevin Y. Yip, Stephen Kwok-Wing Tsui, Qin Cao

## Abstract

Large language models (LLMs) can learn new tasks by in-context learning (ICL), but it is unknown whether this ability reliably transfers to biological sequence classification. Here, we systematically evaluate how demonstration selection, shot count, and prompting strategies affect performance across 20 general-purpose LLMs. Using antibody characterization as a representative test case, we compare zero-shot, few-shot, and chain-of-thought (CoT) ICL on three classification tasks: humanness, antigen specificity, and isotype. Our results reveal a clear performance hierarchy: while zero-shot prompting performs near chance, few-shot prompting with randomly selected demonstrations improves performance, showing that LLMs can perform ICL using biological sequences from minimal supervision. However, matching protein-language model (pLM)-based classifier accuracy is only achieved when using label-diverse demonstrations drawn from antibodies similar to the query sequence. To leverage this insight, we introduce Sim-ICL, a framework that automatically retrieves such demonstrations. Using only 32-shot prompting, Sim-ICL achieves performance competitive with pLM-based classifiers, matching or outperforming them in two of the three tasks. Furthermore, reasoning-oriented prompts yield marginal gains and often produce fluent but biologically incorrect rationales, suggesting that current CoT explanations function as after-the-fact rationalizations rather than capturing mechanistic determinants of antibody properties. From these experiments, we derive practical design principles for ICL on biological sequences: use similarity-based, label-diverse demonstrations and modest shot counts, and treat reasoning prompts primarily as post hoc narratives rather than drivers of performance. Sim-ICL implements these principles in a streamlined, prompt-based framework for antibody sequence classification and, in principle, could be adapted to other biological sequence tasks.

## Introduction

In-context learning (ICL) allows large language models (LLMs) to perform new tasks using information in the prompt, without any task-specific parameter updates^1^. ICL is typically categorized into two major approaches: zero-shot learning, which provides no demonstrations, and few-shot learning, which incorporates a few demonstrations (i.e., “shots”) before asking the LLM to make predictions^1^. In contrast to ICL, fine-tuning updates an LLM’s parameters using thousands to hundreds of thousands of labeled training instances, often achieving strong performance at the cost of a large, task-specific dataset^1^. Interestingly, few-shot ICL can sometimes match or even outperform fine-tuned models on language tasks while using only a handful of labeled demonstrations^1,2^. However, it remains unclear whether these ICL capabilities transfer to biological sequence classification tasks, where inputs are raw amino acid or nucleotide sequences rather than natural language.

ICL has demonstrated initial success beyond natural language-related tasks, including chemistry^3–5^, genomics^6,7^, and single-cell cell-type annotation^8^ tasks. In chemistry, ICL has shown competitive performance on tasks such as molecule property classification and design^5^. However, in bioinformatics, most applications still treat text as the primary input, using LLMs for genomic information retrieval, including retrieving gene genomic location^6^ and nomenclature^7^ tasks, and biomedical text processing^2^, rather than directly using biological sequences for classification. Furthermore, rather than utilizing LLMs for direct classification, recent studies employ them primarily as feature extractors. In these frameworks, LLMs are prompted to generate textual descriptions of genes^9^ or drugs^10^, which are subsequently converted to embeddings to train machine learning (ML) models. Here, we use general-purpose LLMs to perform direct in-context sequence classification using raw amino acid sequences. To the best of our knowledge, this setting remains underexplored.

Applying ICL to biological sequences can support antibody characterization^11^ and accelerate both therapeutic and vaccine development^12^. Antibody characteristics, including antigen binding affinity, specificity, immunogenicity, and developability, are ultimately assessed through *in vivo* evaluations. These evaluations involve efficacy and side effect profiling in animal models and patients, which are time-consuming and laborious processes^12,13^. Computational methods are therefore widely used to prioritize candidates^12–15^. Initially, ML models were hindered by a reliance on structural data, which is often scarce^12^. More recently, ML models trained on large high-throughput sequencing datasets have demonstrated that sequence-only representations can effectively predict antibody properties, such as immunogenicity^16,17^, specificity^18^, and binding affinity^19^, often using protein language model (pLM) embeddings as features.

These advances raise a natural question: Can LLMs, using ICL alone, leverage biological sequence data to achieve performance comparable to specialized sequence-based ML models, and how should prompts be designed to enable this? Here, we address this question using antibody sequence data as a case study, evaluating ICL as a practical alternative to task-specific models and identifying the prompt design principles needed for robust performance. Rather than proposing a new architecture, we systematically investigate the potential and limitations of ICL, analyzing which design choices, including demonstration selection, number of shots, sequence representation, and model choice, most strongly affect performance.

In this work, we introduce Sim-ICL, a sequence-similarity-based prompting framework tailored for users with minimal coding experience to perform biological sequence classification, and systematically derive practical prompt design principles from our experiments. Sim-ICL uses only sequence-derived features to retrieve similar labeled demonstrations and construct few-shot prompts, which makes the approach, in principle, applicable beyond antibodies to other biological sequence tasks. The resulting design rules for choosing demonstrations, setting shot counts, and deciding when reasoning prompts help or hurt extend to sequence-based prediction problems beyond antibodies. We evaluate it using 20 general-purpose LLMs from five families, including GPT^1,20^, Claude^21^, Gemini^22^, LLaMA^23,24^, and Mistral^25,26^. When we use Sim-ICL to select in-context demonstrations, few-shot ICL achieves performance comparable to supervised ML models trained on large, labeled datasets with state-of-the-art pLM embeddings. Also, we assess the reproducibility and stability of these results across models and prompting conditions. Finally, we provide an open-source implementation, including code and prompt templates, to facilitate reuse in other biological sequence-classification settings.

## Results

### Study overview

ICL has transformed natural language processing, yet its potential for biological sequence analysis remains relatively unexplored. Here, we systematically evaluated 20 representative LLMs across five families (including GPT, Claude, Gemini, LLaMA, and Mistral) **(Fig. 1a, Supplementary Table 1, 2, and Methods)**. These selected LLMs have demonstrated strong performance in ICL across multiple fields^1,5^ and achieved high rankings on LLM leaderboards^27,28^.

**Fig. 1.**
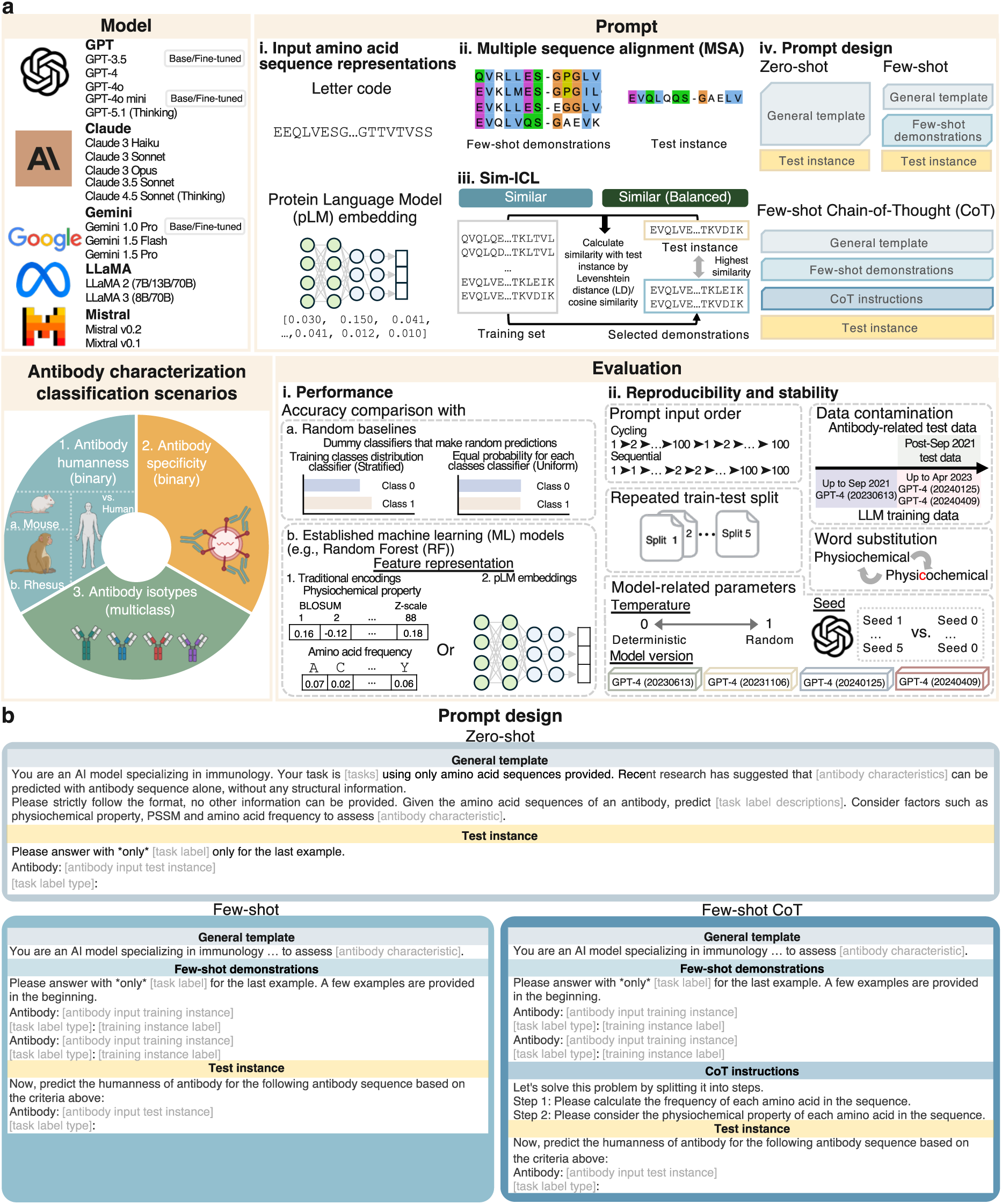
Study overview and prompt design. In this work, we apply ICL to three antibody characterization scenarios by 20 LLMs from five different families. **a** Schematic overview of Sim-ICL and benchmark. Figure created with BioRender.com released under a Creative Commons Attribution-NonCommercial-NoDerivs 4.0 International license. **b** Illustration of ICL prompt design. Detailed prompt designs are given in the **Supplementary Fig. 1-3**.

We designed our benchmarks around three practically motivated antibody characterization scenarios **(Fig. 1a)** that capture core properties relevant to therapeutic design and analysis: humanness, antigen specificity, and heavy-chain isotype. These scenarios represent fundamental antibody characteristics and have been introduced in prior work^12,29^. Scenario 1 (humanness) is a binary classification task motivated by immunogenicity, as more human-like antibodies typically elicit fewer immune responses^16,30^. To cover a range of complexity, we further split this into distinguishing human vs. mouse sequences (Scenario 1a) and human vs. rhesus sequences (Scenario 1b, which provides a more realistic humanness benchmark due to higher sequence similarity). Scenario 2 (specificity) tests whether sequences bind a defined antigen (SARS-CoV-2), a long-standing challenge in computational antibody design^15,31^ and vaccine development^32^. Scenario 3 (isotype) is a multiclass classification problem over heavy-chain isotypes, representing a basic but widely used antibody characteristic^29^. Dataset construction and preprocessing are detailed in **Methods** and **Supplementary Table *3***.

We evaluated ICL using different sequence representations (letter codes vs. pLM embeddings) and input formats (with or without multiple sequence alignment [MSA]). We introduced Sim-ICL, a sequence-similarity-based prompting framework for few-shot demonstration selection utilizing Levenshtein distance (LD) **(Methods)**. We also designed different ICL prompt settings (zero-shot, few-shot, and few-shot Chain-of-Thought [CoT]) **(Fig. 1a, b, and Supplementary Fig. 1-3).** To objectively quantify ICL’s performance, we compared it against random baselines and established ML models, particularly state-of-the-art models utilizing pLM embeddings for feature encoding **(Fig. 1a).** Additionally, we combined ICL with fine-tuning to evaluate its impact on performance **(Fig. 1a and Methods)**, and systematically evaluated the reproducibility and stability of the ICL results **(Fig. 1a)**. Throughout the study, each prompt was repeated five times to evaluate the consistency of the results.

Before comparing ICL to established ML models across all three scenarios, we first investigated a fundamental question: under what prompting conditions does ICL achieve effective performance on antibody sequences? Given that few-shot ICL keeps the model parameters fixed and infers a decision rule from the limited labeled demonstrations in the prompt, the selection of demonstrations is critical. We therefore began by comparing three basic settings: zero-shot prompts without demonstrations, few-shot prompts with randomly selected demonstrations, and few-shot prompts with demonstrations selected using sequence similarity (Sim-ICL), using the easiest task, humanness classification between human and mouse sequences (Scenario 1a).

### Similarity-based demonstrations are essential for robust sequence ICL

Selecting appropriate demonstrations is a key factor in the success of few-shot ICL. Studies have shown that selecting demonstrations similar to test instances can improve few-shot ICL’s performance in language-related^33^ and chemistry^3,5^ tasks. Existing antibody characterization ML models often rely on sequence similarity as an indicator of clonal relatedness for prediction^34,35^. Motivated by this, we introduced Sim-ICL for biological sequence classification in few-shot ICL and compared its performance to zero-shot and the default few-shot setting with randomly selected demonstrations. Specifically, we evaluated six strategies that differed in whether demonstrations were chosen at random or by similarity (Sim-ICL), and whether the order and balance of positive and negative labels were controlled **(Fig. 2a and Methods)**. Balanced selection mitigates the majority-label bias (e.g., all demonstrations being positive instances) that arises when selecting demonstrations based solely on similarity, which can lead to a lack of diversity in the ground-truth labels of the demonstrations **(Supplementary Fig. 4)**.

**Fig. 2.**
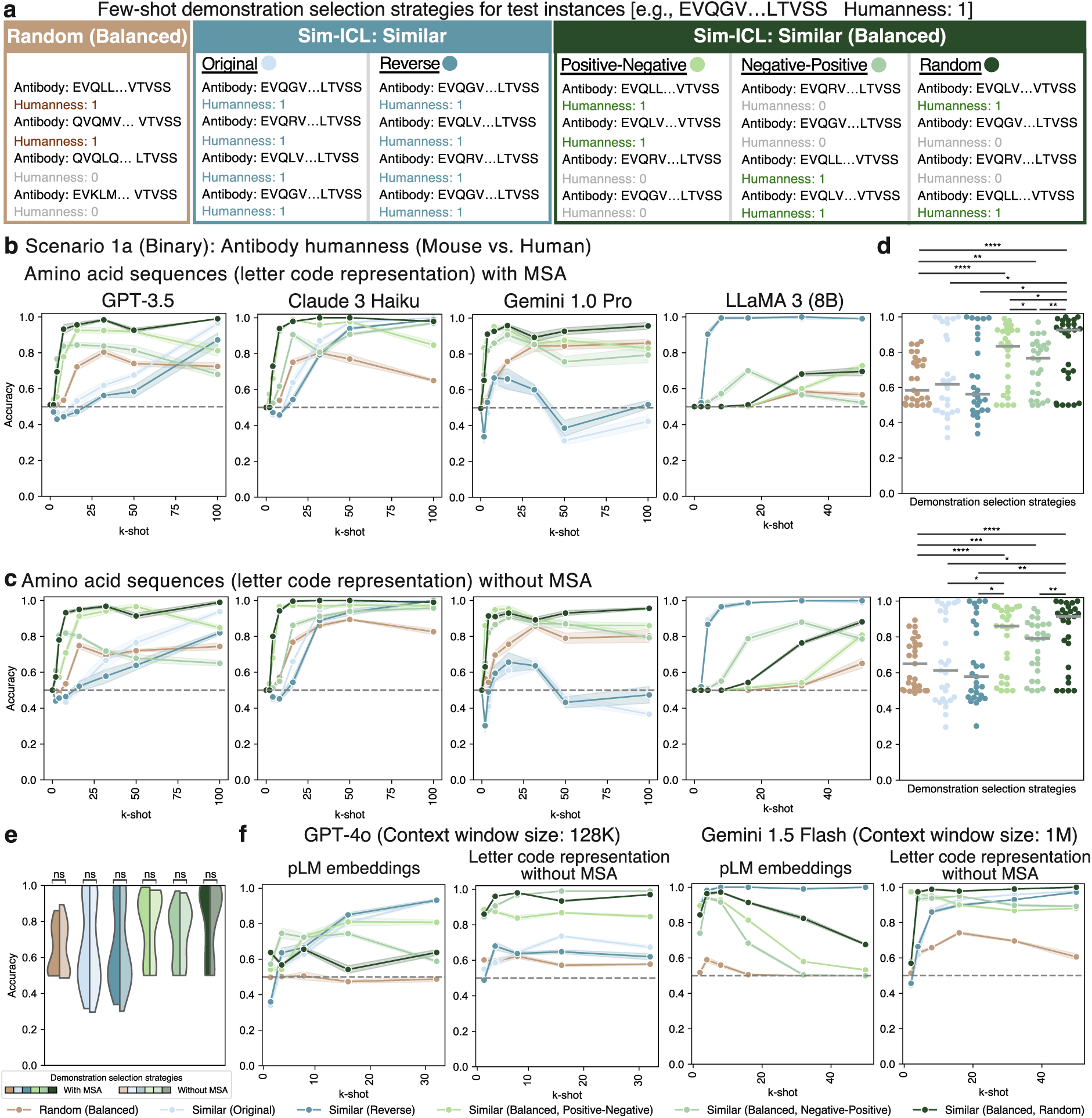
Accuracy evaluation of zero-shot and few-shot demonstration selection strategies using different sequence representations. **a** Illustration of few-shot demonstration selection strategies for a test instance. **b,c** Zero-shot and few-shot (2-, 4-, 8-, 16-, 32-, 50-, 100-shot) ICL accuracy with amino acid sequence input (**b**) with and (**c**) without MSA using different demonstration selection strategies in affordable LLMs. The shaded area indicates standard deviation of five repeated predictions. The dashed line indicates the random baseline trained with full training set **(Methods)**. **d** Statistical test between the demonstration selection strategies of all four evaluated LLMs. Each dot represents an average accuracy of a LLM with a *k*-shot prompt. The grey lines indicate the median for all observations in each demonstration selection strategy. **e** Statistical test between the performance of MSA and without MSA input for each demonstration selection strategy. The darker color represents accuracy with MSA, while the paler color represents accuracy without MSA. **f** Few-shot ICL accuracy using pLM embeddings as input. *P* values in **c** and **d** are based on two-sided paired Wilcoxon rank-sum test: ns: *P* > 0.05; \**P* ≤ 0.05; \*\**P* ≤ 0.01; \*\*\**P* ≤ 0.001; \*\*\*\**P* ≤ 0.0001.

Similarity-based demonstrations dramatically improved humanness classification on an easy benchmark (human vs. mouse), whereas zero-shot prompting failed. Specifically, we evaluated six demonstration strategies in Scenario 1a **(Fig. 1a**), using low-cost LLMs from the four families. In this setting, LLM performance depended critically on both the demonstration selection strategy and the number of in-context demonstrations. All models failed with zero-shot prompting **(Fig. 2b, c)**, indicating that naïve prompting and pre-training alone are insufficient to achieve robust performance for antibody classification tasks. Instead, effective ICL requires carefully selected labeled demonstrations. Accuracy improved with increasing *k* (i.e., more demonstrations) **(Fig. 2b, c)**. At optimal *k*, Sim-ICL significantly outperformed the random strategy **(Fig. 2d)**, enabling near-perfect accuracy **(Fig. 2b, c, and Supplementary Fig. 5)**. The optimal strategy differed by model: GPT-3.5, Claude 3 Haiku, and Gemini 1.0 Pro performed best with Similar (Balanced) selection, while LLaMA 3 (8B) achieved peak performance using the Similar strategy **(Fig. 2b, c, and Supplementary Fig. 5)**.

Next, we tested whether Sim-ICL remains effective when demonstrations lack label diversity. Since the Similar strategy can occasionally retrieve demonstrations that all share the same label **(Supplementary Fig. 4)**, we investigated whether performance would degrade if the prompt contained only positive demonstrations. To test this, we constructed single-label prompts using only positive instances from the Similar (Balanced) strategy. Strikingly, these positive-only prompts reduced performance to random levels across all four low-cost models, whereas GPT-4 remained above chance **(Supplementary Fig. 6)**. In every setting, prompts containing both positive and negative labels consistently outperformed their positive-only counterparts. Together, these results demonstrate that both similarity-based retrieval and label diversity are critical for effective antibody humanness classification. This suggests a general design principle for sequence-based ICL: prompts should include demonstrations that are sequence-similar to the query while strictly maintaining label diversity.

### Effective few-shot sequence ICL does not rely on MSA

To determine if the evolutionary context provided by MSAs improves prediction, we evaluated ICL performance using unaligned sequences (without MSA). Previous evidence suggested that using MSAs instead of unaligned amino acid sequences improved performance in antibody humanness classification by deep learning models^16^. We therefore repeated the above evaluations using unaligned amino acid sequences to assess if this conclusion applies to few-shot ICL **(Supplementary Fig. 7)**.

The same low-cost LLMs showed highly similar performance trends, irrespective of the use of MSA **(Fig. 2b, c)**. The difference in accuracy between the with-MSA and without-MSA formats was not statistically significant across all settings **(Fig. 2e)**. This mirrors findings in pLMs, where MSA-free pLMs showed comparable performance to MSA-based pLMs on several protein classification tasks^36^. These results suggest that for ICL on biological sequences, simple unaligned amino acid representations may suffice, reducing the need for computationally expensive MSA preprocessing. For simplicity, we therefore used sequences without MSA in our subsequent analyses.

### Long-context LLMs can treat pLM embeddings as in-context numeric features

As pLM embeddings are widely used features in antibody characterization tasks^29^, we asked whether LLMs can also effectively use pLM embeddings as an alternative to raw amino acid letter codes. In a proof-of-concept experiment, raw letter code representations were replaced with AntiBERTy embeddings (**Methods)**, and cosine similarity was used instead of LD to select similar demonstrations. To accommodate the larger token footprint of embedding-based prompts, we focused on two economical long-context models, GPT-4o and Gemini 1.5 Flash.

Similarity-based prompting enabled LLMs to predict antibody humanness from pLM embeddings with high accuracy **(Fig. 2f)**. In Gemini 1.5 Flash, the pLM embedding representation achieved comparable accuracy to the letter code representation under the Similar strategy **(Fig. 2f),** and reached near-perfect accuracy with as few as four shots. However, unlike letter-code representations that improved monotonically with k, increasing k did not always produce better results in long-context LLMs under the Similar (Balanced) strategy. This observation is consistent with previous studies in language-related tasks^37^. These findings underscore the versatility of LLMs in leveraging different forms of representation; further advances in long-context LLMs may better exploit pLM embeddings.

### Sim-ICL is competitive with specialized pLM-based ML models in diverse biological scenarios

We next investigated whether Sim-ICL could extend effectively to three clinically relevant antibody characterization scenarios of increasing difficulty (Scenario 1b, Scenario 2, and Scenario 3 in **(Fig. 1a)**). These scenarios exhibit higher sequence overlap and more subtle class differences than the basic human-mouse scenario (Scenario 1a) **(Supplementary Fig. 8)**. To evaluate performance in these more realistic settings, 20 general-purpose LLMs were assessed using 16- and 32-shot similarity-based prompts and compared directly to established ML models trained on full datasets leveraging AntiBERTy pLM embeddings^5^ as features **(Methods)**.

Consistent with our initial findings on the simpler scenario **(Fig. 2b, c)**, few-shot, task-specific demonstrations remained necessary in all evaluated scenarios. Zero-shot prompting **(Fig. 1b, Supplementary Fig. 1, 2, and Methods)** failed to exceed chance-level performance across the three complex antibody characterization scenarios, with accuracies close to random guessing for all evaluated LLMs. In contrast, few-shot ICL with either random or similarity-based demonstration selection strategies significantly outperformed zero-shot ICL **(Supplementary Fig. 9 and Supplementary Table 4b)**. These results indicate pre-trained knowledge alone is insufficient for antibody characterization and that task-specific demonstrations are required for effective LLM performance on these challenging non-natural language tasks^1^.

Having established the necessity of few-shot demonstrations, we next evaluated the few-shot ICL performance in each scenario and compared it with established ML models.

**Scenario 1b: Humanness (human vs. rhesus).** We first assessed antibody humanness using human and rhesus heavy chain sequences. Because rhesus sequences are genetically closer to humans than mice are^16^, this scenario provides a more realistic humanness benchmark than the human-mouse task in Scenario 1a **(Supplementary Fig. 8a, b)**. Similarity-based selection strategies, including the Similar and the Similar (Balanced) strategies, consistently yielded the best performance across all evaluated LLMs **(Fig. 3a)**. All but four models (GPT-4o mini and the three LLaMA 2 variants) outperformed both traditional encoding-based (amino acid frequency and physiochemical encodings) methods **(Methods),** with most achieving near-perfect accuracy (>0.9) **(Fig. 3a)**. Notably, three LLMs (Claude 3.5 Sonnet, LLaMA 3 (8B), and Mistral v0.2) attained a perfect accuracy of 1.0, matching the state-of-the-art AntiBERTy^38^-based pLM classifier **(Fig. 3a)**.

**Fig. 3.**
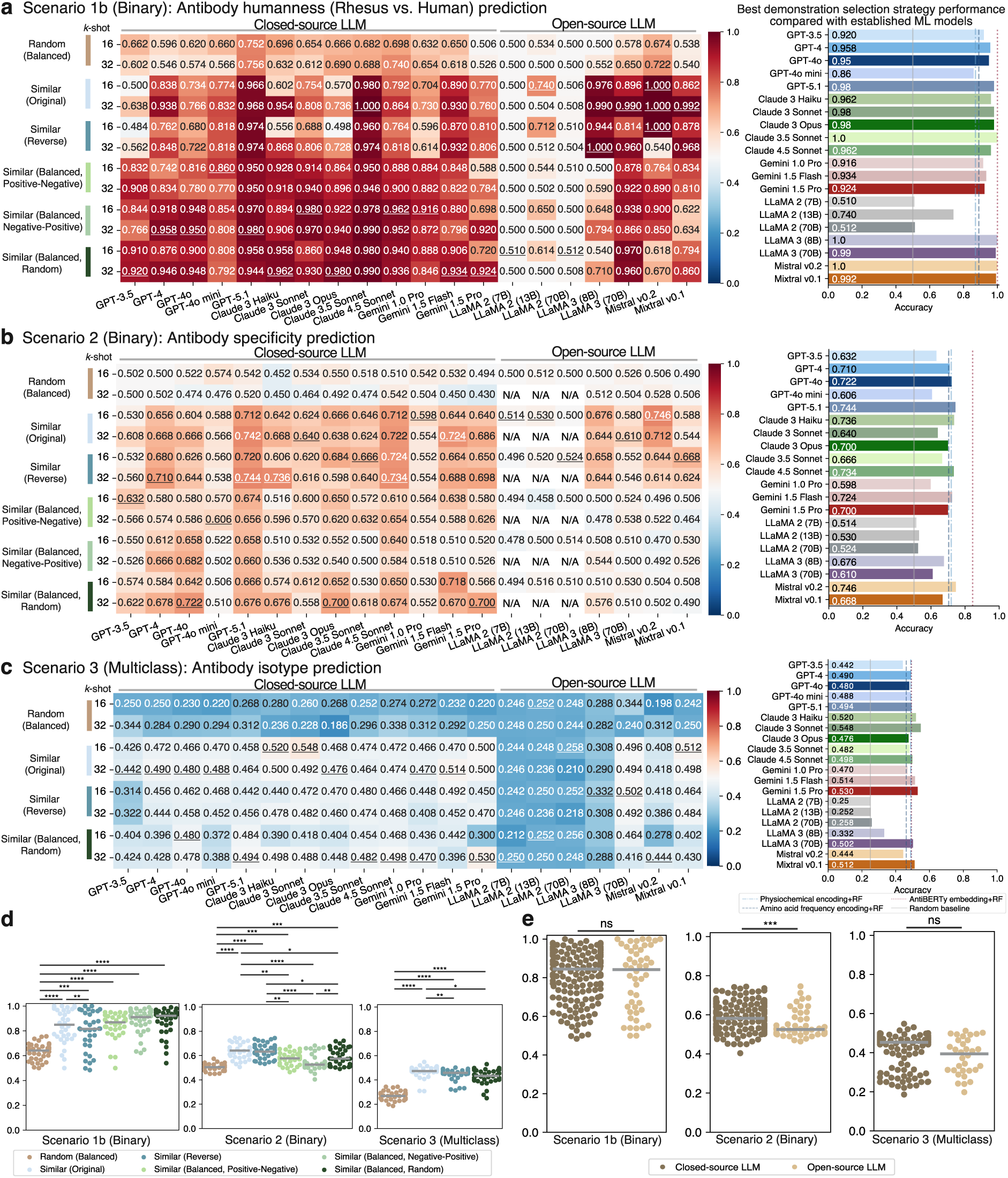
Few-shot ICL accuracy in three antibody characterization classification scenarios. Evaluation of few-shot (16- and 32-shot) using different demonstration selection strategies with 20 LLMs in three antibody characterization classification scenarios. **a-c** Few-shot ICL accuracy and its best demonstration selection strategy accuracy compared with established ML models across 20 LLMs in (**a**) Scenario 1b, (**b**) Scenario 2, and (**c**) Scenario 3. Each entry shows the average accuracy of five repeated predictions. Underlined values indicate the best performance for that LLM. **d,e** Statistical test (**d**) between the demonstration selection strategies and (**e**) between closed-source and open-source models. Each dot represents the average accuracy of an LLM with a k-shot prompt. Note that LLaMA 2 models were excluded in all the comparison in **d** and **e** due to their underperformance compared to other models. P values in **d** are based on two-sided paired Wilcoxon rank-sum test and in **e** are based on two-sided Mann-Whitney U test: ns: P > 0.05; *P ≤ 0.05; **P ≤ 0.01; ***P ≤ 0.001; ****P ≤ 0.0001.

**Scenario 2: Specificity.** The second binary task distinguished SARS-CoV-2 binding from non-binding antibodies. This scenario proved more difficult due to strong class overlap in Uniform Manifold Approximation and Projection (UMAP) space **(Supplementary Fig. 8c)**. Four LLMs (GPT-4o, Claude 3 Haiku, Gemini 1.5 Flash, and Mistral v0.2) outperformed traditional encoding-based baselines (accuracy ≈ 0.71). However, the AntiBERTy-based pLM classifier remained strongest overall (accuracy = 0.845), while the best few-shot ICL performance reached 0.746 (Mistral v0.2) **(Fig. 3b)**. Consistent with Scenario 1b, similarity-based selection yielded the strongest results among ICL strategies **(Fig. 3b)**.

**Scenario 3: Isotype.** The third scenario involved multiclass isotype classification, a challenging task reflected by poor class separation **(Supplementary Fig. 8d)** and modest performance of established ML models **(Fig. 3c)**. Impressively, six LLMs (Claude 3 Haiku, Claude 3 Sonnet, Gemini 1.5 Flash, Gemini 1.5 Pro, LLaMA 3 [70B], and Mixtral v0.1) outperformed the AntiBERTy-based pLM classifier (accuracy=0.492) **(Fig. 3c)**. The same pattern held here, where similarity-based selection again outperformed random prompting **(Fig. 3c)**.

Overall, similarity-based demonstration selection strategies consistently outperformed random selection across all three scenarios **(Fig. 3d)**. Within this framework, the 32-shot setting yielded the highest accuracies for the majority of models (Scenario 1b: 13 out of 20 LLMs, 65.0%; Scenario 2: 11 out of 17 LLMs, 64.7%; Scenario 3: 12 out of 20 LLMs, 60.0%) **(Fig. 3a-c)**.

To understand whether these gains reflect ICL behavior rather than a simple majority-label shortcut, we compared LLM predictions under the Similar strategy with those of a majority-vote classifier (**Methods**). Across LLMs, we observed heterogeneous behaviour: LLaMA 3 (70B) showed strong alignment with the majority-vote baseline, whereas other models, such as GPT-3.5, Claude 3 Sonnet, and Gemini 1.5 Flash, displayed substantially lower alignment scores **(Supplementary Fig. 10)**. When we extended this analysis to the ground-truth labels, the limitations of the majority-vote baseline became clear: although it can reach perfect accuracy (accuracy=1.0) in highly skewed, relatively simple tasks such as Scenario 1b, its accuracy drops to 0.29 in more challenging settings (Scenario 3), indicating that its performance collapses in complex settings **(Supplementary Fig. 10)**. In contrast, the best-performing LLM using the Similar strategy maintain higher accuracy and lower variance across scenarios. This advantage persists even when the demonstration labels are balanced in the Similar (Balanced) strategy, where the majority-label shortcut is unavailable **(Fig. 3a-c)**. Together, these results show that similarity-based few-shot ICL is not only more stable but also outperforms the simple majority-vote heuristic and that its gains are not a result of a majority-label shortcut.

No single LLM or model family dominated performance across all the scenarios, consistent with previous studies on biomedical^2^ and long-context text processing^37^ tasks. Open-source LLMs performed comparably to closed-source LLMs in two (Scenario 1b and 3) of the three evaluated scenarios **(Fig. 3e)**. Interestingly, within the same LLM family (e.g., Claude 3), the most advanced model (i.e., Claude 3 Opus) did not always produce the best performance **(Fig. 3b, c)**. Overall, our results show that using 32 similarity-based demonstrations from Sim-ICL yields strong and reliable few-shot ICL performance across all antibody characterization scenarios, with performance comparable to established ML models in two of the three scenarios. For subsequent analyses, we therefore use a 32-shot Similar (Balanced, Random) strategy and exclude the underperforming LLaMA 2 models.

### Ineffectiveness of reasoning prompts in antibody property prediction and interpretation

Motivated by the success of CoT prompting in arithmetic and logical reasoning tasks^39,40^, we tested three CoT variants for antibody classification: (i) CoT steps only, (ii) LLM-generated answers, and (iii) manually corrected answers **(Methods)**. All three variants consistently underperformed simpler non-CoT prompts across all scenarios. Manual inspection of the reasoning traces generated by CoT revealed systematic calculation errors, particularly in amino acid frequency counts, which is consistent with recent reports^41,42^ **(Supplementary Fig. 11, 12, and Methods)**. Paradoxically, models performed even worse when we replaced their own incorrect calculations with manually corrected answers **(Supplementary Fig. 12, 13)**. Among the three CoT variants, the “CoT steps only” strategy performed best and was therefore used in subsequent analyses **(Supplementary Fig. 13)**. However, even this best-performing variant still underperformed simpler non-CoT prompts across all three evaluation scenarios **(Fig. 4a and Supplementary Table 4c)**. This indicates that the explicit reasoning steps tested here did not improve antibody-related predictions, in contrast to prior gains reported for math- and logic-focused tasks.

**Fig. 4.**
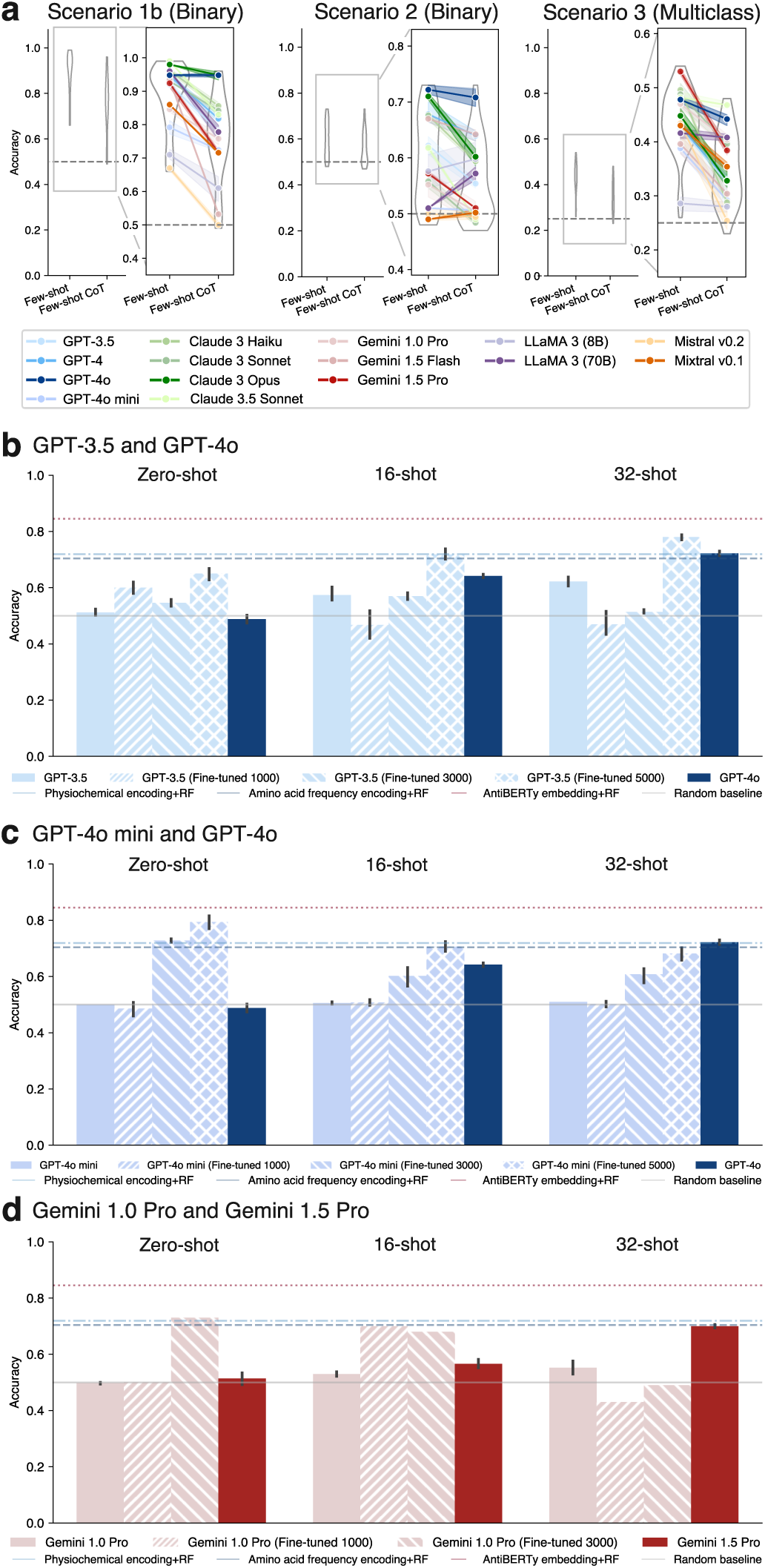
ICL prompt accuracy evaluation in base, fine-tuned, and advanced models. **a** Accuracy of few-shot (32-shot) and few-shot CoT (32-shot) prompts in three antibody characterization scenarios. The dashed line indicates the random baseline trained with full training set. **b-d** Performance comparison of zero-shot and few-shot (16- and 32-shot) prompts of an LLM in its base, fine-tuned, and advanced models in Scenario 2: **b** GPT-3.5’s base model, fine-tuned models, and advanced model (i.e., GPT-4o); **c** GPT-4o mini’s base model, fine-tuned models, and advanced model (i.e., GPT-4o); and **d** Gemini 1.0 Pro’s base model, fine-tuned models, and advanced model (i.e., Gemini 1.5 Pro) in Scenario 2. The number in each fine-tuned model’s name denotes the number of training instances used in fine-tuning.

Given that explicit reasoning steps did not improve prediction accuracy, we next investigated whether the generated reasoning traces could at least provide biologically meaningful explanations. Qualitative inspection of outputs from reasoning models (i.e., GPT-5.1 [Thinking], Claude 4.5 Sonnet [Thinking]), which autonomously generate internal reasoning chains without explicit prompting, showed fluent, domain-relevant narratives that referenced concepts such as CDR3 composition, constant regions, and structural motifs **(Supplementary Fig. 14)**. However, manual inspection revealed that these traces frequently misapplied core immunological concepts. For instance, models often erroneously identified variable-region sequences as constant-region “IGHM signatures” or treated CDR3 patterns as isotype-specific markers. These findings indicate that such reasoning traces primarily reflect post hoc rationalizations of model outputs rather than mechanistic accounts of antibody biology.

### Moderate-scale fine-tuning combined with ICL approaches pLM-based ML performance

We investigated whether fine-tuning on medium-sized datasets could substantially improve LLM performance on the difficult antibody specificity task and, when combined with few-shot ICL, approach the accuracy of the pLM-based (AntiBERTy-based) ML model. Unlike ICL, which keeps model parameters fixed, fine-tuning requires additional compute and labeled data in exchange for potentially higher task-specific accuracy^1^. To quantify this trade-off, we fine-tuned three LLMs with different architectures: GPT-3.5, GPT-4o mini, and Gemini 1.0 Pro on moderate-sized training sets (i.e., 1,000, 3,000, and 5,000 instances) from Scenario 2 **(Supplementary Fig. 15 and Methods)**. Performance was then compared to the corresponding base and more advanced models (GPT-4o and Gemini 1.5 Pro) under zero-shot and few-shot ICL prompts.

In the zero-shot setting, fine-tuned models generally outperformed both their base and advanced counterparts, with fine-tuned GPT-4o mini achieving the best LLM accuracy (0.794) **(Fig. 3b and 4b-d)** and further narrowing the gap to the pLM-based ML model (accuracy=0.845) **(Fig. 4b-d)**. In the 16-shot setting, all fine-tuned Gemini 1.0 Pro models outperformed their base and advanced models, whereas only GPT-3.5 and GPT-4o mini fine-tuned with 5,000 training instances exceeded their base and advanced models **(Fig. 4b-d)**. At higher shot counts, performance depended on architecture: a fine-tuned GPT-3.5 model with a 32-shot prompt (accuracy=0.780) outperformed ML models using traditional encodings **(Fig. 4b and Methods)**. In contrast, fine-tuned Gemini 1.0 Pro models showed diminished performance in 32-shot ICL, underperforming both their base and advanced models **(Fig. 4d)**. Together, these results show that fine-tuning on medium-sized datasets, combined with few-shot prompting, improves LLM performance on difficult specificity tasks. This middle-ground approach comes close to, but does not consistently exceed, the accuracy of the pLM-based ML model.

### Reproducibility and stability evaluation

Several factors and hyperparameters may influence the performance of ICL. To ensure the reproducibility and stability of our results, we conducted a comprehensive investigation into these factors. We chose GPT models as representatives for this analysis due to their widespread popularity.

#### Effect of prompt input order

Prompt input order had a negligible impact on ICL performance. Comparing cycling versus sequential input ordering **(Fig. 1a and Methods)** revealed similar accuracies across models **(Fig. 5a)**, indicating that LLM responses are robust to the choice of input order.

**Fig. 5.**
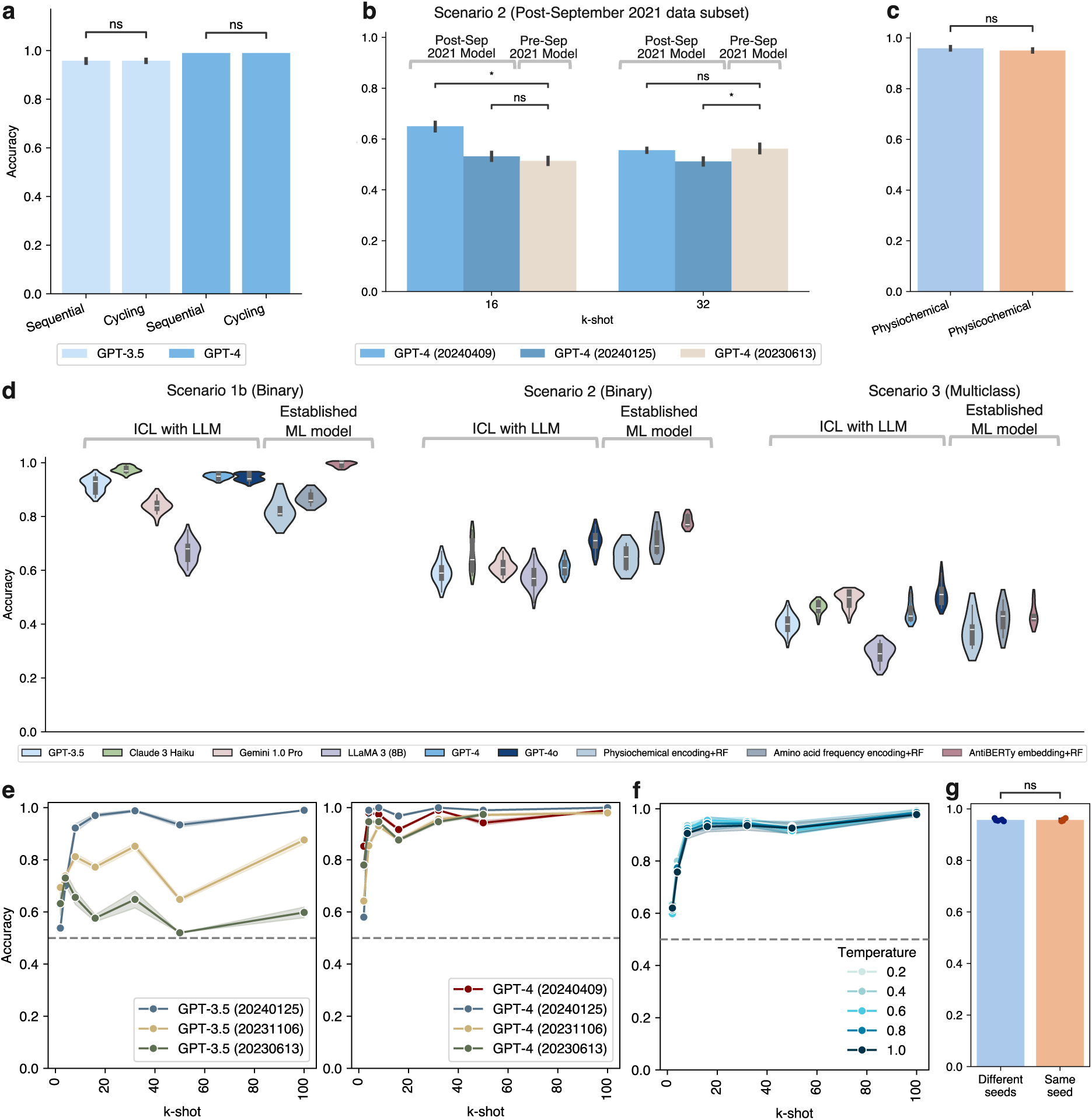
Reproducibility and stability evaluation. **a** Accuracy comparison between cycling and sequential prompt input order. **b** Data contamination comparison in LLMs with training data cut-off in September 2021 (i.e., pre-September 2021 model: GPT-4 [20230613]) and December 2023 (i.e., post-September 2021 models: GPT-4 [20240125] and GPT-4 [20240409]) with post-September 2021 data as the test set. **c** Alternative spelling performance comparison with the word changed from “physiochemical” to “physicochemical”. **d** Accuracy evaluation with existing ML models using five train-test split for evaluating ICL robustness. **e** Accuracy evaluation for GPT model variations. **f** Accuracy evaluation for parameter *temperature*. The dashed line indicates the random baseline trained with full training set in **e** and **f**. **g** Accuracy comparison for parameter *seed*. Each dot represents the mean accuracy of five repeated responses per *seed*. *P* values in **a-c** and **g** are based on two-sided Mann-Whitney *U* test: ns: *P* > 0.05; \**P* ≤ 0.05.

#### Assessment of data contamination

Data contamination, which occurs when LLMs are exposed to the test set during pre-training, may inflate performance^43^. If data contamination exists, pre-trained LLMs that potentially encounter the test set will be expected to show better performance compared to the models trained solely on the data before the test set was published **(Methods)**. Following Li et al.^43^, we selected SARS-CoV-2 antibody specificity (Scenario 2) data published in year 2022 (post-September 2021) and applied it to LLMs with training data cut-off in September 2021, i.e., pre-September 2021 model (GPT-4 [20230613]), and training data cut-off in December 2023, i.e., post-September 2021 models (GPT-4 [20240125] and GPT-4 [20240409]), to test whether data contamination exists.

In the 16-shot setting, despite the statistically significant differences between pre- and one of the post-September 2021 models (GPT-4 [20240409]), we observed that the other post-September 2021 model showed a statistically insignificant difference (GPT-4 [20240125]). Furthermore, in the 32-shot setting, the post-September 2021 models either exhibited statistically insignificant differences or even significantly worse performance compared to the pre-September 2021 model **(Fig. 5b)**. Together, these observations indicate that our results were unlikely to be influenced by data contamination.

#### Tolerance to alternative terminology

We examined ICL’s tolerance for alternative spellings or misspellings. Both words “physiochemical”^18^ and “physicochemical”^29^ were widely used to describe antibody characteristics in the literature **(Methods)**. The word choices did not affect the predictions **(Fig. 5c)**. Our results suggest that ICL exhibits some tolerance to alternative terms.

#### Robustness across train-test split

To address the potential issues associated with using a small test set, such as overfitting to specific examples, insufficient coverage of data variability, and increased variance in performance metrics, we performed multiple train-test splits using five random seeds. This ensured the model was tested on diverse data subsets. No instances were shared between different test sets. The evaluation was expanded to also include Claude 3 Haiku, Gemini 1.0 Pro, and LLaMA 3 (8B) for a more comprehensive analysis. The ICL’s performance trend in different LLMs was consistent with that in previous sections **(Fig. 5d)**. Also, ICL’s performance variability was comparable to established methods using traditional encodings and pLM embeddings **(Fig. 5d)**. This indicates that ICL is robust to data variability.

#### Model-related hyperparameters Effect of model version

Next, we evaluated the impact of model-related hyperparameters on prediction stability **(Methods)**. GPT-3.5 showed greater variation, while GPT-4 remained relatively stable across different versions. Surprisingly, the latest model did not always perform better, with GPT-4 (20240125) outperforming GPT-4 (20240409) in all the evaluated k-shot settings except 2-shot **(Fig. 5e)**. Researchers need to take extra caution when selecting different versions of LLMs, as the latest LLMs may not always yield the best results.

#### Effect of temperature parameter

The *temperature* parameter, which controls randomness and creativity in LLM responses, did not affect predictions in few-shot ICL in our scenario **(Fig. 5f, Supplementary Fig. 16, and Methods)**.

#### Effect of random seed

Given the non-deterministic nature of LLMs’ responses, we assessed the impact of the seed parameter on output consistency **(Methods)**. The performance using different seeds was not significantly different compared to that using the same seed **(Fig. 5g)**. This suggests that the seed parameter of the OpenAI application programming interface (API) has a minimal effect on the output consistency, and that the model’s performance is relatively stable across different random initializations.

## Discussion

In our benchmarks, general-purpose LLMs matched or outperformed pLM-based ML models on antibody prediction in two of three evaluated scenarios (Scenarios 1 and 3) when Sim-ICL was applied. However, the pLM-based ML model retained the advantage on the hardest task, antigen specificity prediction (Scenario 2). Zero-shot prompts stayed near chance on all three tasks, indicating that pretraining on natural language alone is insufficient for our antibody prediction tasks. In contrast, few-shot prompting with randomly selected demonstrations already improved performance relative to zero-shot baselines, showing that ICL with task-specific demonstrations is effective. Sim-ICL extended these gains further, approaching state-of-the-art accuracy by adding label-diverse sequences similar to the query. Finally, CoT prompting did not improve accuracy, and as the explanations from reasoning models were often biologically implausible, we interpret them mainly as post hoc narratives rather than the true basis for the models’ decisions.

We distill these findings into four practical design principles for applying ICL to biological sequence classification. First, demonstrations should be curated to be both sequence-similar to the query and label-balanced to prevent majority-class bias. Second, simple unaligned amino acid sequences should be adopted as the default input representation, as this minimizes preprocessing without sacrificing accuracy compared to MSAs or current pLM embeddings. Third, prioritize using a small, well-chosen set of demonstrations over using exhaustive shot counts. We recommend allocating approximately 32 demonstrations, as this modest number already achieves competitive performance with pLM-based classifiers in our benchmarks, with additional shots yielding diminishing returns. Finally, for current LLMs, CoT reasoning should be treated as an optional mechanism for post hoc interpretability rather than a primary tool for optimizing predictive performance.

These principles naturally organize antibody sequence modeling into three complementary approaches, each suited to different resource constraints. First, pure ICL offers a rapid, prompt-based option for low-data settings when similar labeled sequences are available, leveraging sequence-similar demonstrations to achieve competitive performance on foundational tasks without any training infrastructure. Second, fine-tuned LLMs occupy a middle ground, utilizing moderate-sized datasets to bridge the accuracy gap in more complex scenarios. Third, pLM-based ML models, which typically require greater programming expertise to implement, remain the top-performing choice in our benchmarks, especially on the challenging specificity prediction task. Together, these approaches define a flexible spectrum allowing researchers to balance coding requirements, model accessibility, and predictive accuracy. Because our ICL framework relies solely on raw sequence inputs and labeled demonstrations, the same design principles can be extended from antibodies to a broad range of biological sequence-classification problems.

These mechanistic insights have immediate practical implications. Existing antibody sequence databases, such as Observed Antibody Space (OAS)^44^, already facilitate similarity searches for a specific sequence of interest, enabling the construction of similarity-based ICL prompts without requiring coding knowledge. Our results therefore point to a concrete workflow in which antibody characterization can be carried out by interacting with an LLM via similarity-based prompts, either directly through a chatbot interface **(Supplementary Fig. 17)** or through user-friendly APIs. Additionally, given that open-source and closed-source LLMs showed comparable ICL performance in the two evaluated scenarios, researchers can prioritize open-source models when they need customization and transparency ^45^. Together, the combination of high performance, prompt-based accessibility, and flexible deployment options makes ICL a practical complement to established ML models for antibody characterization tasks.

Nevertheless, our study has certain limitations. First, the test set sizes are relatively small **(Methods)** due to the high inference costs associated with LLMs. To mitigate potential robustness concerns, we have repeated the experiments across multiple train-test splits and observed stable results. We anticipate that future LLM advances will lower expenses, enabling more extensive testing. Second, our evaluation of similarity was primarily binary, contrasting high-similarity demonstrations with random selection. We did not systematically quantify the specific similarity thresholds required for success. Future work should quantitatively evaluate how varying degrees of sequence similarity (e.g., 90%, 70%, or 50%) impact performance to determine the precise similarity thresholds at which ICL can outperform established ML models. We also anticipate that future LLM advancements will go beyond reliance on sequence similarity, potentially capturing intrinsic biological features to enable accurate predictions even when similar demonstrations are scarce. Third, we only explored a small set of CoT prompt designs, which underperformed non-CoT prompts. More tailored reasoning prompts might yield benefits in antibody-related tasks^46^. Finally, we have focused on antibodies as a representative case. Future work should test whether these design principles generalize to other sequence modalities, such as enzymes or genomic regulatory elements. We anticipate that further theoretical investigations into ICL^47^, alongside empirical performance assessments, will validate the efficacy of ICL in regression applications, such as binding affinity prediction, and in generative applications, like therapeutic antibody design.

## Methods

### Dataset collection and processing

The datasets used for each antibody characterization scenario, including the number of instances in each target class for both training and test sets, were summarized in **Supplementary Table 3**. We ensured that all instances were unique (i.e., no duplicate instances) within each scenario.

#### Scenario 1: Antibody humanness prediction

##### Sequences with MSA

The MSA files containing the aligned antibody heavy chain sequences were directly downloaded from Ramon et al.^16^. This downloaded dataset had an equal number of sequence instances across all the target classes, resulting in a balanced set. According to Ramon et al.^16^, the source of human, mouse and rhesus antibody heavy chain sequences were obtained from the OAS^44^ database. The retrieved sequences were aligned using the AHo^48^ numbering scheme, with ANARCI^49^ software, which numbers each residue based on its structural role, resulting in aligned sequences of length 149. There were 20,000 unique sequences for the dataset used in Scenario 1a and 1b, respectively.

In MSA, the numbering of amino acids is based on their structural roles rather than their original positions in the sequence^49^. This numbering scheme provides a pre-defined “static position template” that generates an identical output format for a specific sequence, regardless of the number or identity of other sequences aligned with it **(Supplementary Fig. 7)**. This ensures the alignment does not rely on other sequences and reflects structural importance, allowing the model to learn structural patterns from the training set without being exposed to position-specific information from the test set.

##### Sequences without MSA

For the sequence input without MSA, we retrieved the sequences by removing the gaps (denoted by “-”) of the downloaded sequences (encoded by amino acid letter code representation) in the MSA files by our custom bash script. This resulted in unaligned amino acid sequence for each antibody from the original MSA.

#### Scenario 2: Antibody specificity prediction

Antibody heavy and light chain sequences from single-cell paired heavy and light chain full-length B-cell receptor (BCR) V(D)J sequencing were downloaded from Wang et al.^29^, covering datasets across ten different studies. Following Wang et al.’s ^29^ suggestion for optimal performance, we concatenated heavy and light chain sequences for antibody specificity prediction. There were 15,538 unique concatenated sequences for the dataset used.

#### Scenario 3: Antibody isotype prediction

Antibody heavy chain sequences were downloaded from the same study used in Scenario 2 (Wang et al.^29^). To enable the learning, classes with fewer than 50 instances were discarded. The target classes for the heavy chain isotype class prediction included IGHM, IGHD, IGHG, and IGHA. After filtering and deduplication, 0.81 million unique heavy chain sequences remained.

##### Train-test split

For each antibody characterization scenario, the full dataset was randomly divided into the training set and test set. Due to cost considerations, we set each test set to contain 100 instances. The test sets were set balanced with equal number of instances per target class **(Supplementary Table 3)**. All few-shot demonstrations were selected exclusively from the training set to ensure no data leakage in the prompts.

### Demonstration selection and prompt design

#### ICL demonstration selection

For the random strategy, i.e., Random (Balanced), an equal number of instances from each target class were randomly selected exclusively from the training set.

#### Sim-ICL

For the amino acid letter code-based representation in the Sim-ICL, including both the Similar and the Similar (Balanced) strategies, sequence similarity was assessed using LD between each training instance and each test instance. For the Similar (Balanced) strategy, an equal number of similar instances were selected from each target class. In *k*-shot learning, we chose the *k* most similar *k* training instances (those with the smallest LD) for each test instance.

Similarly, for the pLM embedding-based representation in the Sim-ICL, *k*-shot demonstrations were chosen from the training set based on the cosine similarity instead of LD. Specifically, we first used the state-of-the-art pre-trained antibody-specific pLM, AntiBERTy^38^, to generate an embedding for each antibody sequence. The resulting embedding for each sequence was then averaged over its length to ensure uniform embedding size across sequences of different lengths, in line with established practices^29,36^. This averaging process was done by the Python script in Wang et al.^29^. Each averaged AntiBERTy embedding (i.e., the pLM embedding) had a dimensionality of 512. Subsequently, *k*-shot demonstrations were selected from the training set based on the highest cosine similarity to the test instance, using the pLM embeddings. To minimize token usage, the pLM embeddings were further rounded to three decimal places and input as vectors in the few-shot prompts. All other prompt structures remained identical to those in the amino acid letter code-based representation.

#### Prompt design

A prompt refers to a set of instructions and context provided as input to guide the language model’s response generation. In most APIs, prompts can be provided to different roles, including “system”, “user”, and “assistant” roles. The “system” role, included at the start of the prompt, provides context, instructions, or other information to prime the model for the specific use case. The “user” role contains the actual user queries. The “assistant” role contains the model’s responses to the user queries, based on the context provided in the “system” role and “user” role. These roles work together, with the “system” message setting the stage, the “user” message providing the actual user queries, and the “assistant” message containing the model’s response to the user queries considering the overall context.

All prompts in this work were input into the “user” role, unless otherwise specified. The prompts were designed following previous publications^3,5^ and prompt engineering guides from OpenAI^50^ and Anthropic^51^. We experimented with different prompt designs, each containing elements such as task descriptions, examples, and reasoning steps. In all the prompts, a new line was represented by the “\n” character.

#### Zero-shot

The zero-shot prompt is designed to evaluate LLMs’ performance based solely on its pre-trained knowledge, without the aid of any task-specific demonstrations. For the zero-shot prompt, we provided a context-setting instruction to the LLM, asking it to act as an immunology expert, following the template shown in **Fig. 1b** and **Supplementary Fig. 2**.

#### Few-shot

To construct the few-shot prompt, we added the few-shot demonstrations before the test instance **(Fig. 1b and Supplementary Fig. 1)**. The entire prompt remained the same as zero-shot prompt except the inclusion of few-shot demonstrations **(Supplementary Fig. 1, 2)**.

#### Few-shot CoT

CoT reasoning steps, based on antibody properties related to antibody characterization, including calculating amino acid frequency and considering amino acid physiochemical properties, suggested by previous literature^39,40,52^, were added to the few-shot prompt. We tested three few-shot CoT prompt structures **(Supplementary Fig. 11)**.

For CoT steps alone, we followed Reynolds et al.^39^ to construct the prompts. We placed the two following reasoning steps after the few-shot demonstrations: “Let’s solve this problem by splitting it into steps.\nStep 1: Please calculate the frequency of each amino acid in the sequence.\nStep 2: Please consider the physiochemical property of each amino acid in the sequence.”. These two reasoning steps were kept consistent across all three CoT prompt structures **(Supplementary Fig. 11a)**.

For CoT with LLM-generated answers, we referenced both Wei et al.^40^ and Reynolds et al.^39^ to construct the prompts. Initially, we provided the first reasoning step, and the LLM generated a response. We then input this response into the “assistant” role along with the second reasoning step into the “user” role. After receiving the LLM’s response, we input it again into the “assistant” role and the remaining part of the prompt into the “user” role **(Supplementary Fig. 11b, c)**.

For CoT with manual answers, we manually corrected the LLM-generated answers, as LLM-generated answers may contain errors, such as miscalculating amino acid frequencies, similar with findings from recent studies^41,42^ **(Supplementary Fig. 12)**. Unlike the CoT approach with LLM-generated answers, where a query may generate a different response at a time, the reasoning steps in this method remained consistent across all the prompts since we provided the same manually curated answers as the input into the “user” role **(Supplementary Fig. 11d)**.

Once the CoT reasoning steps were complete, the test instances were fed into the “user” role in prompts to generate the final response, using the same prompt structure as before **(Supplementary Fig. 11)**. CoT steps alone were used in the subsequent comparison with zero-shot and few-shot settings **(Fig. 4)** as it showed the best performance among the three CoT variations in in **Supplementary Fig. 13**.

### LLMs

Our selected 20 LLMs cover models of different architectures, with different parameter sizes, and from different organizations, including both closed-source and open-source models **(Supplementary Table 1)**. If an unexpected API interruption occurs (e.g., network interruption), the entire prediction process will restart from the beginning. All the LLMs used a *temperature* of 0.2 (except GPT-5.1 [Thinking] and Claude 4.5 Sonnet [Thinking]), and all the other API parameters were set to default unless specified. For each test instance in each setting, we ran its prompt five times to evaluate the consistency of the results. Each LLM’s response was summarized in **Supplementary Table 4**. The majority of the LLMs (16 out of 20 LLMs, 80.0%) provided formatted responses without any additional words beyond the predicted labels **(Supplementary Table 4 and Methods)**.

The details for running each LLM are described as follows:

**OpenAI (GPT-3.5, GPT-4.0, GPT-4o, GPT-4o mini, and GPT-5.1)**

The prompt was used to query GPT-3.5, GPT-4o, GPT-4.0, and GPT-4o mini via the OpenAI API (v1.11.0), following the official OpenAI guidelines^53^. GPT-5.1 was queried in batch mode via the OpenAI Responses API (v2.8.0), with the reasoning-summary (“thinking summary”) option enabled (using the reasoning parameter). For the OpenAI API, the n parameter is available for generating output n times using a single input prompt. This parameter n was used instead of running a single prompt n times, reducing the number of API calls required. We set n to be 5.

#### Anthropic (Claude 3 Haiku, Claude 3 Sonnet, Claude 3 Opus, Claude 3.5 Sonnet, Claude 4.5 Sonnet)

The prompt was used to query Claude 3 Haiku, Claude 3 Sonnet, Claude 3 Opus, and Claude 3.5 Sonnet via the Anthropic API (v0.21.3) following the official Anthropic guidelines^54^. Claude 4.5 Sonnet was queried via the Anthropic Messages API (v0.73.0) with extended thinking enabled and summarized thinking output using the thinking parameter. Only the final text responses were used in our analyses.

#### Google (Gemini 1.0 Pro, Gemini 1.5 Flash, and Gemini 1.5 Pro)

The prompt was used to query Gemini 1.0 Pro, Gemini 1.5 Flash, and Gemini 1.5 Pro via the Google Generative API (v0.5.3) following the official Google guidelines^55^.

#### Meta (LLaMA 2 and LLaMA 3)

For LLaMA 2, the chat models were used instead of the instruct models as the latter failed to provide appropriate answers **(Supplementary Fig. 18)**. For LLaMA 3, the instruct models were used. For Scenario 1a, it should be noted that LLaMA 3 was chosen over LLaMA2 due to the latter’s limited context window size **(Supplementary Table 1)**.

The [BOS] (beginning of a sentence) and [EOS] (end of a sentence) special tokens were added to the prompt following the prompt format in the Meta official guidelines^56,57^. As LLaMA 3 instruct models were used, [INST] and [/INST] instruction tokens were also added to the prompts for all LLaMA 3 models. For LLaMA 2 (7B) and LLaMA 3 (8B) models, we ran the experiments both locally through Python Hugging Face pipeline (for Scenario 1b) and via the Replicate API (for Scenario 2 and 3), where other models, namely LLaMA 2 (13B and 70B) and LLaMA 3 (70B), solely via the Replicate API (v0.25.2). We transitioned from Hugging Face pipeline to Replicate API for its faster inference speed in our study. Locally, we generated the response by inputting the prompts to the corresponding models (meta-llama/Llama-2-7b-chat-hf and meta-llama/Meta-Llama-3-8B-Instruct) in the Python Hugging Face transformer (v.4.25.1) library on NVIDIA A100 (40G). All the parameters were set based on the official Meta tutorial^58,59^ and the GitHub llama-recipes repository. The *num_return_sequences* parameter is available in the Hugging Face pipeline for generating output multiple times using a single input prompt. For the Replicate API, the formatted prompt was used to query each model to obtain the response.

#### Mistral AI (Mistral and Mixtral)

We formatted prompts according to the Mistral AI guidelines^60,61^, adding the <s> (BOS), [INST] and [/INST] tokens to the Mistral (v0.2) prompt, and the same tokens plus an additional</s> (EOS) token to the Mixtral (8×7b) (v0.1) prompt. Mistral was used both locally via the Python Hugging Face pipeline (for Scenario 1b and 3) and via the Replicate API (for Scenario 2), whereas Mixtral was used solely via the Replicate API. We transitioned from Hugging Face pipeline to Replicate API for its faster inference speed in our study. Locally, we generated the response by inputting the prompt to the corresponding model (mistralai/Mistral-7B-Instruct-v0.2) in the Python Hugging Face transformer (v.4.25.1) library on NVIDIA A100 (40G). The *num_return_sequences* parameter was used in the Hugging Face pipeline to generate output multiple times from a single input prompt. For the Replicate API (v0.25.2), the corresponding formatted prompt was used to query Mistral and Mixtral to obtain the response, respectively.

### Model fine-tuning

#### Fine-tuning prompts

The fine-tuning prompts were created by randomly selecting 1,000, 3,000, or 5,000 instances from the training set. The fine-tuning prompts with 5,000 instances were excluded in Gemini 1.0 Pro due to its API file size limitation. The fine-tuning prompts are illustrated in **Supplementary Fig. 15**.

The details for fine-tuning each LLM are described as follows:

**GPT-3.5 and GPT-4o mini**

The fine-tuning prompts were formatted according to the OpenAI API guidelines, using the JavaScript Object Notation (JSON) structure: {“messages”: [{“role”: “user”, “content”: input_prompt}, {“role”: “assistant”, “content”: “label”}]} **(Supplementary Fig. 15b)**. The resulting JSON files were uploaded to OpenAI servers to create fine-tuned models using the OpenAI API (v1.11.0). Zero-shot and few-shot prompts were then input to the resulting fine-tuned models for evaluation.

#### Gemini 1.0 Pro

The fine-tuning prompts were formatted into Comma-Separated Values (CSV) files with column headers “text_input” and “output” added, following the Gemini guidelines **(Supplementary Fig. 15c)**. The resulting CSV files were uploaded to Google AI Studio web interface for fine-tuning. Zero-shot and few-shot prompts were then input to the resulting fine-tuned models for evaluation.

### Performance evaluation

#### LLM response conversion

The majority of the 20 LLMs directly outputted formatted responses with the predicted label. We referred formatted response as response without any additional information other than the answer (e.g., “1”). To convert the responses into predicted labels, we used the following criteria: across all the scenarios, a predicted label of 1 was assigned to formatted responses of “1”, “1 \n”, “1\n” or “*1*”. For unformatted responses **(Supplementary Table *4*)**, specific phrases were extracted for conversion. In Scenario 1, a predicted label of 1 was assigned to responses containing “Humanness: 1” or “Answer: 1”. In Scenario 2, a predicted label of 1 was assigned to responses containing “Binding: 1” or “Answer: 1” or containing both “1” and “bind”. In Scenario 3, a predicted label of 1 was assigned to responses containing “IGHD” or “Answer: 1”. Other labels in Scenario 3 followed the same rationale. The remaining responses were classified as having a predicted label of 0 in all the scenarios.

#### Evaluation metric

After converting the responses into labels, accuracy was calculated as the evaluation metric by comparing the predictions with the true labels of the test set. Each experiment was repeated five times, and the accuracy was averaged across these five repetitions, with the final average accuracy being reported.

#### Statistical tests

In all the sections involving statistical tests, unless otherwise specified, we used the non-parametric two-sided paired Wilcoxon signed-rank test to compare distributions of continuous values. Mann-Whitney *U* test was used for unpaired performance comparisons between closed-source and open-source models and in the “Assessing factors influencing ICL performance” section. All *p*-values for multiple comparisons were corrected by the Bonferroni correction.

#### Performance comparison between zero-shot ICL and few-shot ICL

The performance comparison is depicted in **Supplementary Fig. 9**. In each scenario, for zero-shot ICL, the final accuracy was averaged over 20 (LLMs) × 5 (repetitions) = 90 accuracy scores. For Scenario 1b, the final accuracy of few-shot ICL was averaged over 20 (LLMs) × 6 (demonstration selection strategies) × 2 (16-shot and 32-shot) × 5 (repetitions) = 1,200 accuracy scores. For Scenario 2, the final accuracy of few-shot ICL was averaged over 20 (LLMs) × 6 (demonstration selection strategies) × 1 (16-shot) × 5 (repetitions) + 17 (LLMs, excluding three LLaMA 2 models) × 6 (demonstration selection strategies) × 1 (32-shot) × 5 (repetitions) = 1,110 accuracy scores. For Scenario 3, the final accuracy of few-shot ICL was averaged over 20 (LLMs) × 4 (demonstration selection strategies) × 2 (16-shot and 32-shot) × 5 (repetitions) = 800 accuracy scores.

#### Comparison with random baselines and established models

We compared ICL with random baselines and established ML models.

#### Established ML models

##### Amino acid frequency encoding and physiochemical encoding

Following the Python script in Wang et al.^29^, amino acid frequency and physiochemical encodings of each antibody sequence were extracted from the Python Biopython (v1.7.0) and peptides (v0.3.2) packages, respectively. The amino acid frequency encoding resulted in a feature vector with a dimensionality of 20, while the physiochemical encoding produced a feature vector with a dimensionality of 88.

##### AntiBERTy (pLM) embeddings

We followed the same procedure for generating AntiBERTy embeddings as described in “ICL demonstration selection”. Specifically, we first used the state-of-the-art pre-trained antibody-specific pLM AntiBERTy^38^ to generate an embedding for each antibody sequence. The resulting embedding for each sequence was then averaged over its length to ensure uniform embedding size across sequences of different lengths, in line with established practices^29,36^. This averaging process was done by the Python script in Wang et al.^29^. Each averaged AntiBERTy embedding (i.e., the pLM embedding we finally used) had a dimensionality of 512.

##### The training and test sets of the established ML models

The full training set **(Supplementary Table *3*)** in each scenario was used for training the established ML models. Our first strategy was to train a model using all the instances in the corresponding full training set. We used the RF classifier (sklearn.ensemble.RandomForestClassifier) from the Python scikit-learn (v1.5.0) package, with 1,000 trees and all other hyperparameters set to their default values. The RF model was then tested on exactly the same test set as ICL.

One concern arose from the nature of LLMs in our work: our LLMs outputted binary predictions based on our prompts, prohibiting us from using threshold-free classification evaluation metrics such as the area under the receiver operating characteristic curve (AUROC). In contrast, classifiers such as RF used in this work initially produce continuous prediction scores that need a threshold to be transformed into binary predicted labels. The class with the highest score becomes the model’s predicted class by default, meaning in binary classification, the default threshold is 0.5. However, it is often overlooked that the choice of an appropriate threshold is influenced by the training class imbalance and can affect the final result of threshold-dependent metrics like accuracy. To minimize this potential impact, we carefully designed our second strategy. Specifically, for Scenario 2 and 3, which had class imbalances, we randomly downsampled the full training set to create a balanced training set and trained the RF model with the balanced set (Scenario 2: each class contained 6,830 instances; Scenario 3: each class contained 43,364 instances). For Scenarios 2 and 3, we repeated the downsampling process ten times, obtained ten different balanced training sets, trained the RF model on these ten balanced sets, and averaged the performance on the test set **(Supplementary Table 5)**.

The RF models using our second strategy achieved better performance **(Supplementary Table 5)**. To construct a more challenging and rigorous comparison, we used the better performance as the performance of the established models **(Fig. 3)**.

##### Random baseline

The theoretical random baseline (i.e., random guessing) yields an accuracy of 0.5 for binary classifications and 0.25 for 4-class classifications due to the class balance of our test set. To empirically confirm the accuracy, we constructed two baselines using the dummy classifier function (sklearn.dummy.DummyClassifier) from the Python scikit-learn (v1.5.0) package. The “stratified” baseline predicts class labels based on the observed frequency of each class in the training set, while the “uniform” baseline assigns equal probability to each class. We adopted the two strategies used in the established models. For the first strategy, we further averaged the predictions over 100 random seeds (defined by the *random_state* hyperparameter of the dummy classifier function). For the second strategy, we followed the same procedure as that of the established models. The empirical accuracy aligned closely with the theoretical random baseline accuracy **(Supplementary Table 5)**; therefore, we used the theoretical random baseline as the random baseline in all experiments.

##### Majority voting classifier for label bias evaluation

To evaluate whether LLMs are likely to degenerate into a majority voting classifier when faced with imbalanced label distributions of the demonstrations in few-shot ICL, we created a classifier that always predicts the most frequent label in the demonstrations and compared its performance to LLMs’. We treated the LLM’s predictions (using Similar [Original] strategy) as the true labels and the majority voting classifier’s predictions as the predicted labels. Higher accuracy indicates greater alignment between the LLM and majority vote predictions, while lower accuracy suggests deviation. This approach offers a quantitative assessment of whether LLMs rely heavily on majority label bias in their decision-making or consider additional information beyond the majority vote in the demonstration labels.

##### Reproducibility and stability evaluation

We refer to “reproducibility” as obtaining identical results using the same data, code, and hyperparameters, and “stability” as the model’s performance not being overly sensitive to changes in hyperparameter values. Unless otherwise specified, the following sections used the Similar (Balanced, Random) strategy for all prompt tests in Scenario 1a without MSA.

##### Prompt input order

We evaluated if LLMs’ responses would be affected by two different input orders: cycling and sequential **(Fig. 1a)**. For example, if we have two test instances, each associated with a corresponding prompt, the cycling input order involves alternating between the first and second prompts, repeating this sequence five times. In contrast, the sequential input order consists of inputting the first prompt five times in a row, followed by the second prompt five times. We applied these two input orders across all test instances in the test set.

##### Data contamination

To check for data contamination, following Li et al.^43^, we selected the data generated by a subset of studies that were published in 2022 from Scenario 2. The same train-test split procedure was applied to this subset as described in the previous section. The selected data may overlap with the training data of GPT-4 (20240409) and GPT-4 (20240125), as both models have a training data cut-off of December 2023. However, it does not overlap with GPT-4 (20230613), which has a training data cut-off in September 2021. The performance of models trained with different training data cut-off was compared.

##### Word substitution test

The prompts containing “physiochemical” were compared to those containing “physicochemical”, while all other words in the prompts remained the same.

#### Model-related hyperparameters

##### Model version

We used Scenario 1a with MSA to evaluate different model versions. The OpenAI API offers multiple versions of the same LLM, which typically vary in terms of training data and specific model settings **(Supplementary Table 2)**. For instance, GPT-4 (20240409) and GPT-4 (20240125) represent two distinct versions of the same LLM.

##### Seed

For the different seed test, five distinct seeds (set using the OpenAI API parameter *seed*) were used, each generating five responses with the parameter *n*=5 (for generating response *n* times), resulting in 25 responses. For comparison, a new seed, different from the previous five, was used five times, generating five responses each time using the same parameter *n*=5, for a total of 25 responses.

##### Temperature

In the temperature test, GPT-3.5 was evaluated with temperature values of 0.2, 0.4, 0.6, 0.8, and 1.0 (set by the OpenAI API parameter *temperature*) to examine how temperature variations affect the model’s output.

## Data availability

All data used in this study can be obtained following the procedure described in the “Dataset collection and processing” part of the Methods section.

## Code availability

All code used in this study is openly available in a GitHub repository at https://github.com/madnessfish/bioseq_icl, including the full analysis pipeline and a Jupyter Notebook to facilitate reproducing the results.

## Acknowledgments

QC is supported by National Natural Science Foundation of China under Award Number 32100515 and CUHK direct grant for research under Award Numbers 2022.080 and 2025.031.

## Contributions

SHF and QC conceived the project. KYY, SKWT, and QC supervised the project. SHF and QC designed the computational experiments and data analyses. SHF prepared the data, implemented the methods, conducted the experiments, and analyzed the results. CM and BSHW independently reproduced parts of the results. SHF, ZZ, RW, CM, BSHW, KYL, CH, JZ, KYY, SKWT, and QC interpreted the results. SHF and QC wrote the manuscript. All authors read and approved the final manuscript.

## Ethics declarations

### Ethics approval and consent to participate

No ethical approval was required for this study. All utilized public datasets were generated by other organizations that obtained ethical approval.

## Supplementary Information

**Supplementary Table 1:**
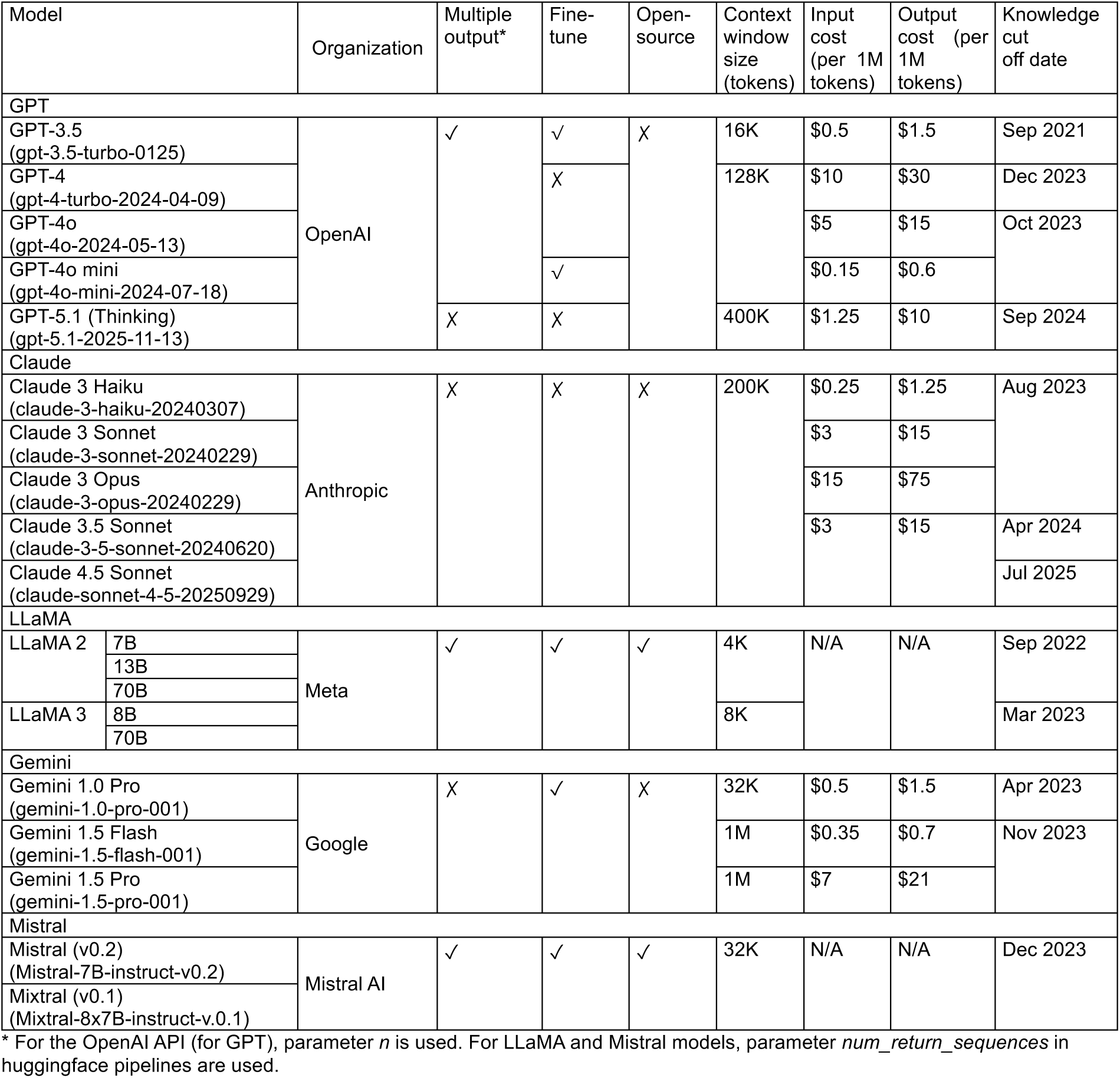
Model and cost summary (Official API, as of 2025/11/24)

**Supplementary Table 2:**
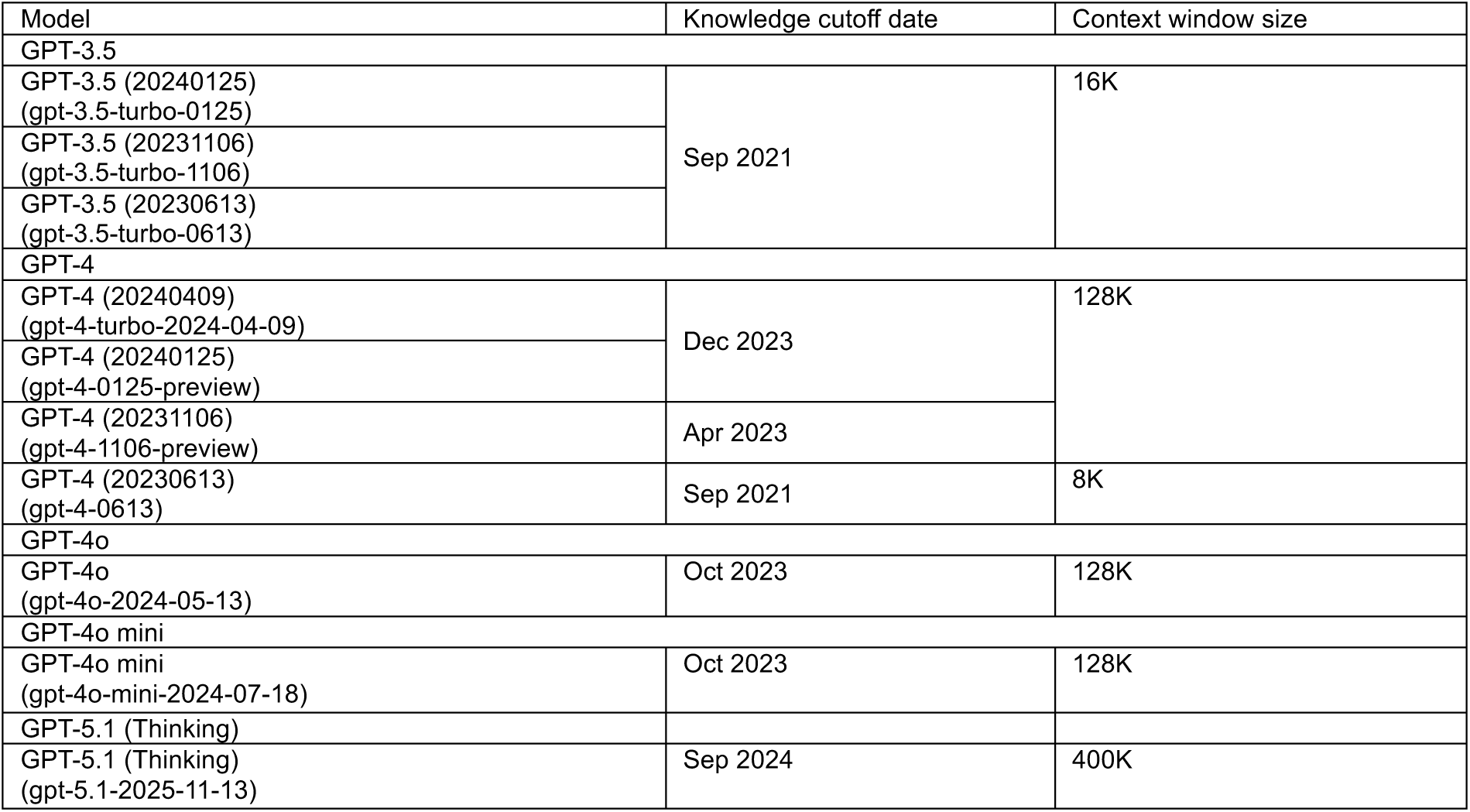
GPT model variations’ knowledge cutoff date and context window size.

| Model | Knowledge cutoff date | Context window size |
| --- | --- | --- |
| GPT-3.5 |  |  |
| GPT-3.5 (20240125)<br>(gpt-3.5-turbo-0125) | Sep 2021 | 16K |
| GPT-3.5 (20231106)<br>(gpt-3.5-turbo-1106) |  |  |
| GPT-3.5 (20230613)<br>(gpt-3.5-turbo-0613) |  |  |
| GPT-4 |  |  |
| GPT-4 (20240409)<br>(gpt-4-turbo-2024-04-09) | Dec 2023 | 128K |
| GPT-4 (20240125)<br>(gpt-4-0125-preview) |  |  |
| GPT-4 (20231106)<br>(gpt-4-1106-preview) | Apr 2023 |  |
| GPT-4 (20230613)<br>(gpt-4-0613) | Sep 2021 | 8K |
| GPT-4o |  |  |
| GPT-4o<br>(gpt-4o-2024-05-13) | Oct 2023 | 128K |
| GPT-4o mini |  |  |
| GPT-4o mini<br>(gpt-4o-mini-2024-07-18) | Oct 2023 | 128K |
| GPT-5.1 (Thinking) |  |  |
| GPT-5.1 (Thinking)<br>(gpt-5.1-2025-11-13) | Sep 2024 | 400K |

**Supplementary Table 3.** Summary of the training and test sets in three antibody characterization scenarios.

| Training set | Number of instances |  |  |  |  |
| --- | --- | --- | --- | --- | --- |
|  | Class 0 | Class 1 | Class 2 | Class 3 | Total number of instances |
| Scenario 1a | 9,950<br>(Mouse) | 9,950<br>(Human) |  |  | 19,900 |
| Scenario 1b | 9,950<br>(Rhesus) | 9,950<br>(Human) |  |  | 19,900 |
| Scenario 2 | 6,830<br>(Non-binder) | 8,608<br>(Binder) |  |  | 15,438 |
| Scenario 3 | 124,947<br>(IGHA) | 43,364<br>(IGHD) | 260,994<br>(IGHG) | 384,573<br>(IGHM) | 813,878 |
| Test set | Number of instances |  |  |  |  |
|  | Class 0 | Class 1 | Class 2 | Class 3 | Total number of instances |
| Scenario 1a | 50<br>(Mouse) | 50<br>(Human) |  |  | 100 |
| Scenario 1b | 50<br>(Rhesus) | 50<br>(Human) |  |  | 100 |
| Scenario 2 | 50<br>(Non-binder) | 50<br>(Binder) |  |  | 100 |
| Scenario 3 | 25<br>(IGHA) | 25<br>(IGHD) | 25<br>(IGHG) | 25<br>(IGHM) | 100 |

**Supplementary Table 4 Summary of LLMs’ response across various prompts and scenarios.** Formatted responses refer to those containing only the answer with no extra information (e.g., “1”, “1\n”). The truncated responses in some LLMs were due to the restricted number of output tokens set by the *max_new_tokens* parameter recommended in each LLM’s user guidelines.

(See the attached PDF file)

**Supplementary Table 5.** Performance of random baselines and established ML models in three antibody characterization scenarios using full and balanced training sets. Note that the original training sets in Scenario 1a and 1b are balanced.

|  | Random baselines |  |  | Established ML models |  |  |
| --- | --- | --- | --- | --- | --- | --- |
|  | Theoretical | Stratified | Uniform | Amino acid frequency encoding + RF | Physiochemical encoding + RF | AntiBERTy + RF |
| Full training set | 0.5 | 0.499 | 0.506 | 0.96 | 0.93 | 1.0 |

Scenario 1b (Binary): Antibody humanness (Rhesus vs. Human) prediction
|  | Random baselines |  |  | Established ML models |  |  |
| --- | --- | --- | --- | --- | --- | --- |
|  | Theoretical | Stratified | Uniform | Amino acid frequency encoding + RF | Physiochemical encoding + RF | AntiBERTy + RF |
| Full training set | 0.5 | 0.499 | 0.506 | 0.89 | 0.87 | 1.0 |

Scenario 2 (Binary): Antibody specificity prediction
|  | Random baselines |  |  | Established ML models |  |  |
| --- | --- | --- | --- | --- | --- | --- |
|  | Theoretical | Stratified | Uniform | Amino acid frequency encoding + RF | Physiochemical encoding + RF | AntiBERTy + RF |
| Full training set | 0.5 | 0.499 | 0.499 | 0.640 | 0.690 | 0.820 |
| Balanced training set |  | 0.502 | 0.494 | 0.709 | 0.719 | 0.845 |

Scenario 3 (Multiclass): Antibody isotype prediction
|  | Random baselines |  |  | Established ML models |  |  |
| --- | --- | --- | --- | --- | --- | --- |
|  | Theoretical | Stratified | Uniform | Amino acid frequency encoding + RF | Physiochemical encoding + RF | AntiBERTy + RF |
| Full training set | 0.25 | 0.251 | 0.244 | 0.45 | 0.43 | 0.46 |
| Balanced training set |  | 0.244 | 0.244 | 0.462 | 0.49 | 0.492 |

**Scenario 1b (Binary): Antibody humanness (Rhesus vs. Human) prediction**

**Supplementary Fig. 1.**
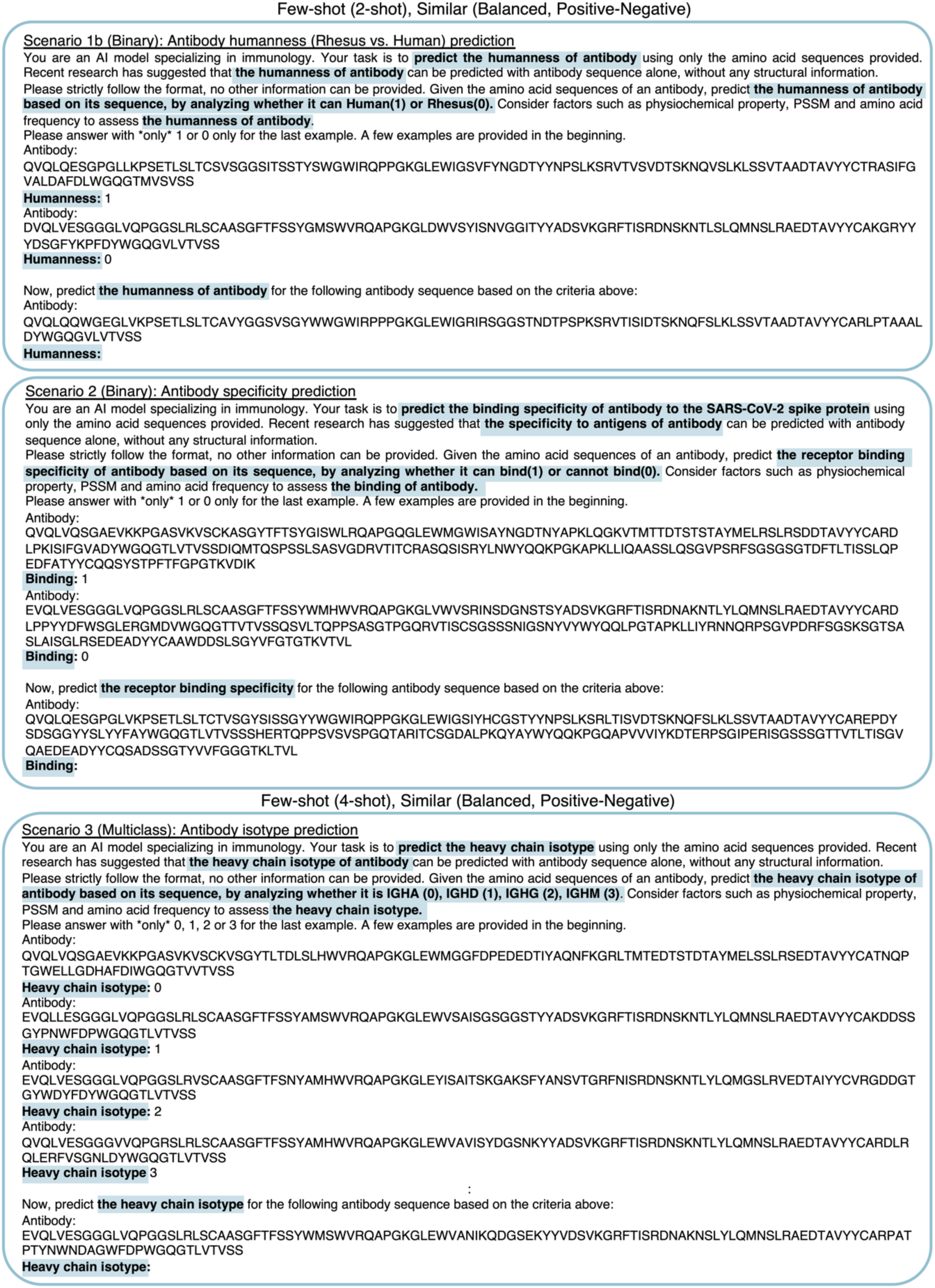
Examples of few-shot prompts in three antibody characterization scenarios. The text highlighted in this figure shows the variable inputs specific to each scenario. All other text remains constant across the examples.

**Supplementary Fig. 2.**
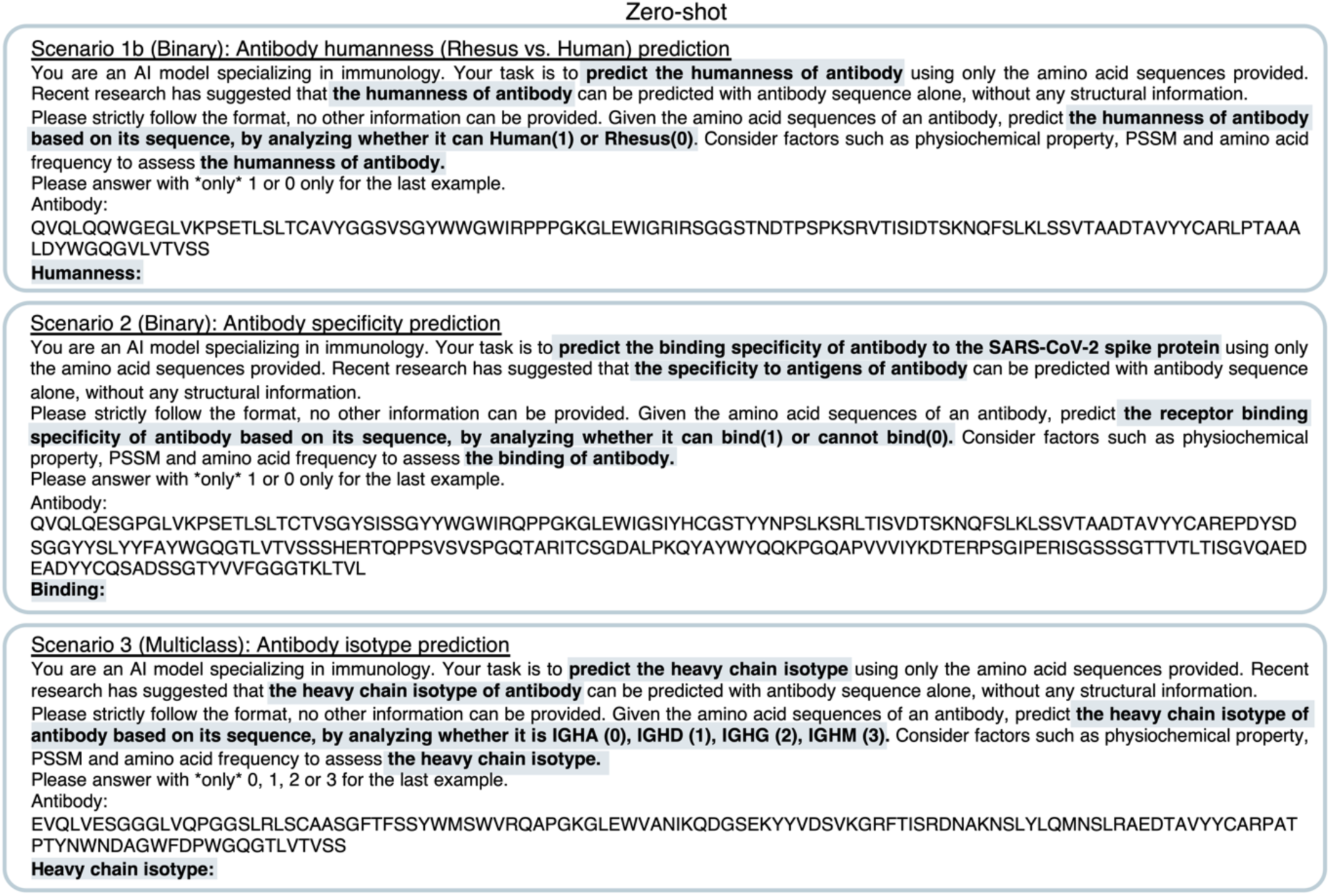
Examples of zero-shot prompts in three antibody characterization scenarios. The text highlighted in this figure shows the variable inputs specific to each scenario. All other text remains constant across the examples.

**Supplementary Fig. 3.**
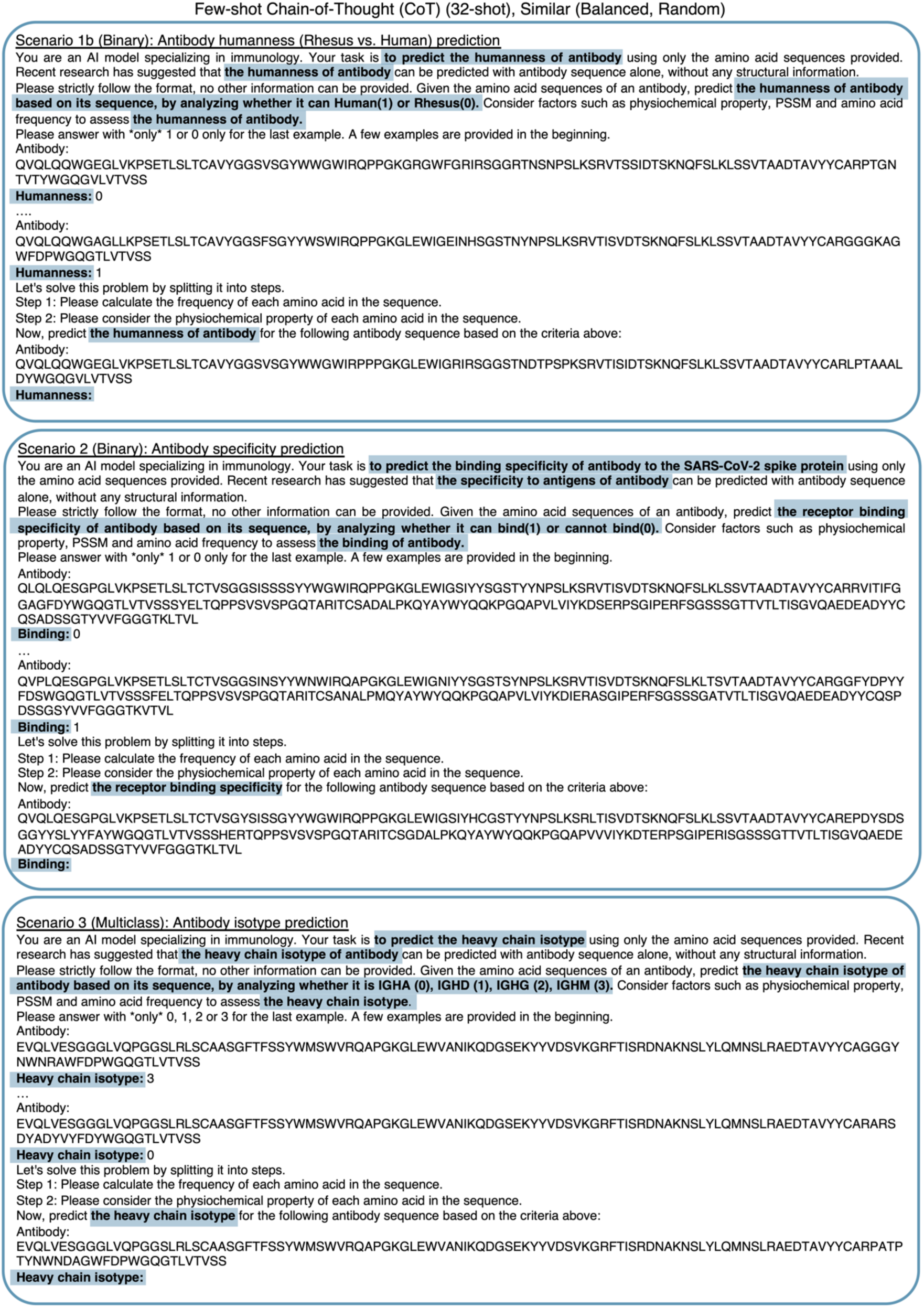
Examples of CoT prompts in three antibody characterization scenarios. The text highlighted in this figure shows the variable inputs specific to each scenario. All other text remains constant across the examples.

**Supplementary Fig. 4.**
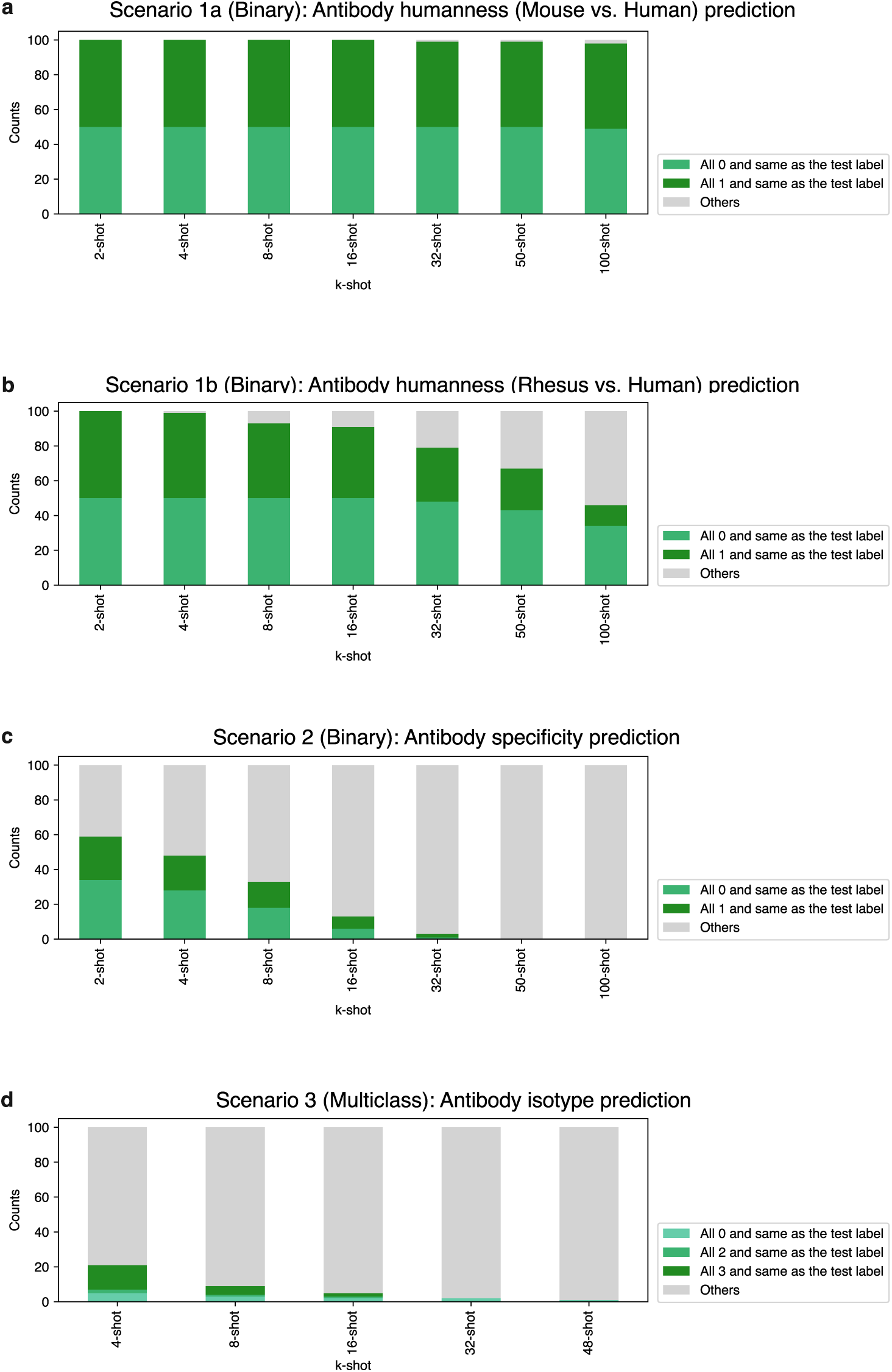
Examination of potential label bias in the Similar strategy in all the evaluated scenarios. Cumulative bar plots showing if all the demonstrations share the same label type (e.g., all the demonstrations have label equals to 1 for a certain test instance) in test prompts (*n*=100) for different *k*-shot settings in Scenario 1a (**a**), Scenario 1b (**b**), Scenario 2 (**c**), and Scenario 3 (**d**). Note that in Scenario 3, there were no cases where all the demonstrations had a label equal to 1.

**Supplementary Fig. 5.**
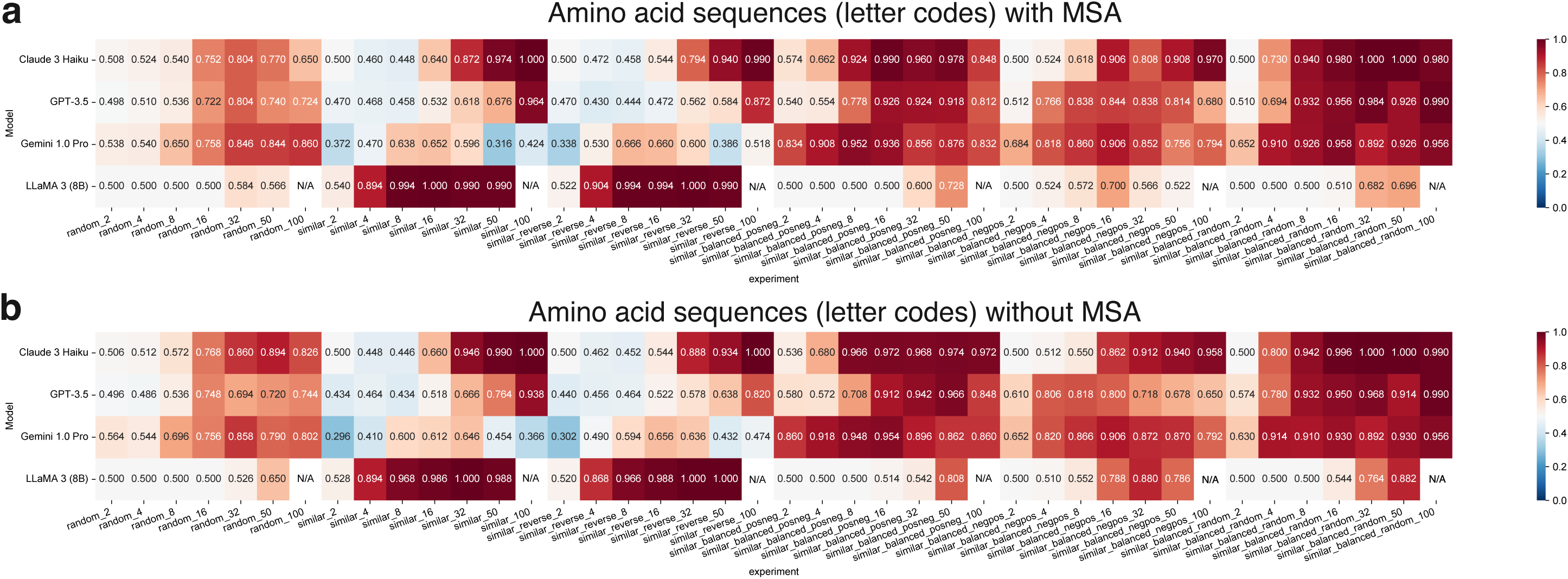
Accuracy of different few-shot demonstration selection strategies in Scenario 1a. The values indicate accuracy for **a** with MSAs and **b** without MSAs.

**Supplementary Fig. 6.**
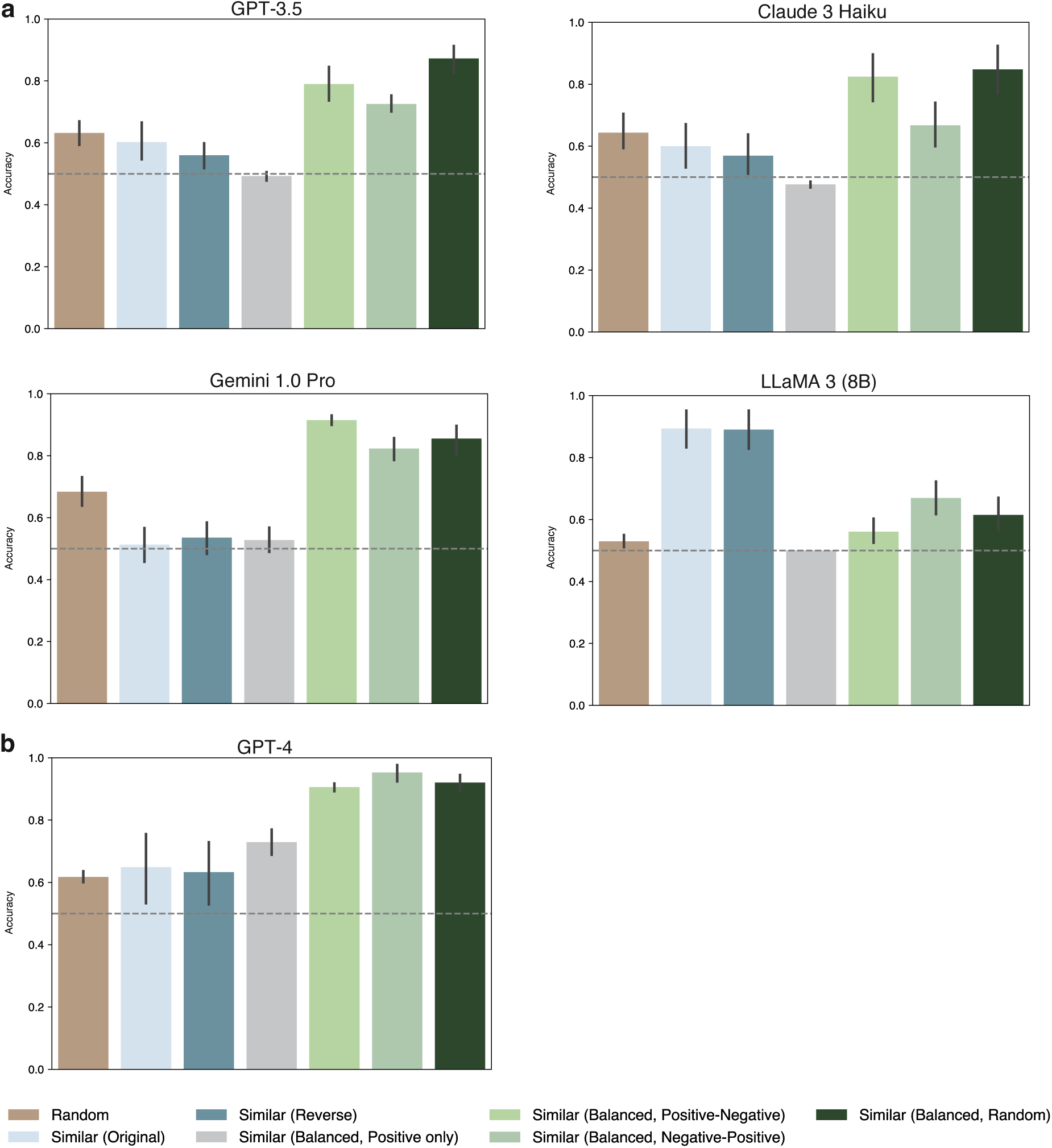
Average few-shot performance for inputting positive labels only in the Similar (Balanced) strategy. Few-shot (including 2-, 4-, 8-, 16-, 32-, 50-, 100-) performance for inputting positive labels only in the Similar (Balanced) strategy in **a** affordable LLMs, including GPT-3.5, Claude 3 Haiku, Gemini 1.0 Pro, and LLaMA 3 (8B), and **b** an advanced model, GPT-4. The dashed line represents the random baseline (accuracy=0.5) of the prediction.

**Supplementary Fig. 7.**
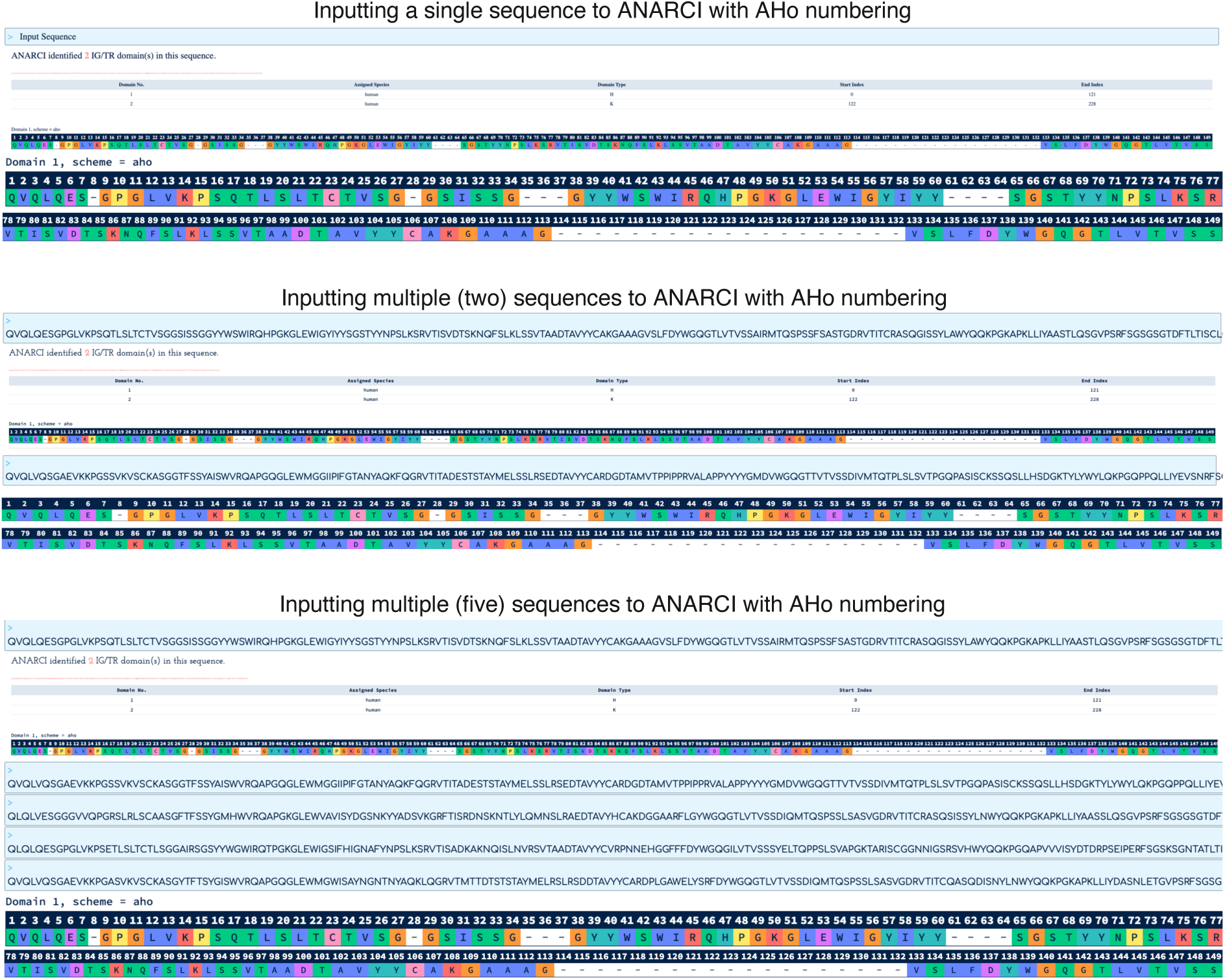
MSA illustrations. The MSAs downloaded from Ramon et al.^1^ contained sequences that were aligned, cleaned, and processed. Non-redundant sequences were aligned to a length of 149 using the widely used ANARCI software and the AHo^2^ numbering scheme. This numbering scheme provides a pre-defined “static position template” that does not rely on specific input sequences. In other words, for a specific sequence, no matter inputting zero, one, two, three, or any numbers of other sequences together with it, the generated output MSA format for this specific sequence is identical. This figure shows an example when using ANARCI software with the AHo numbering scheme, inputting single or multiple sequences produces the same sequence representation, indicating the MSA is independent of other sequences.

**Supplementary Fig. 8.**
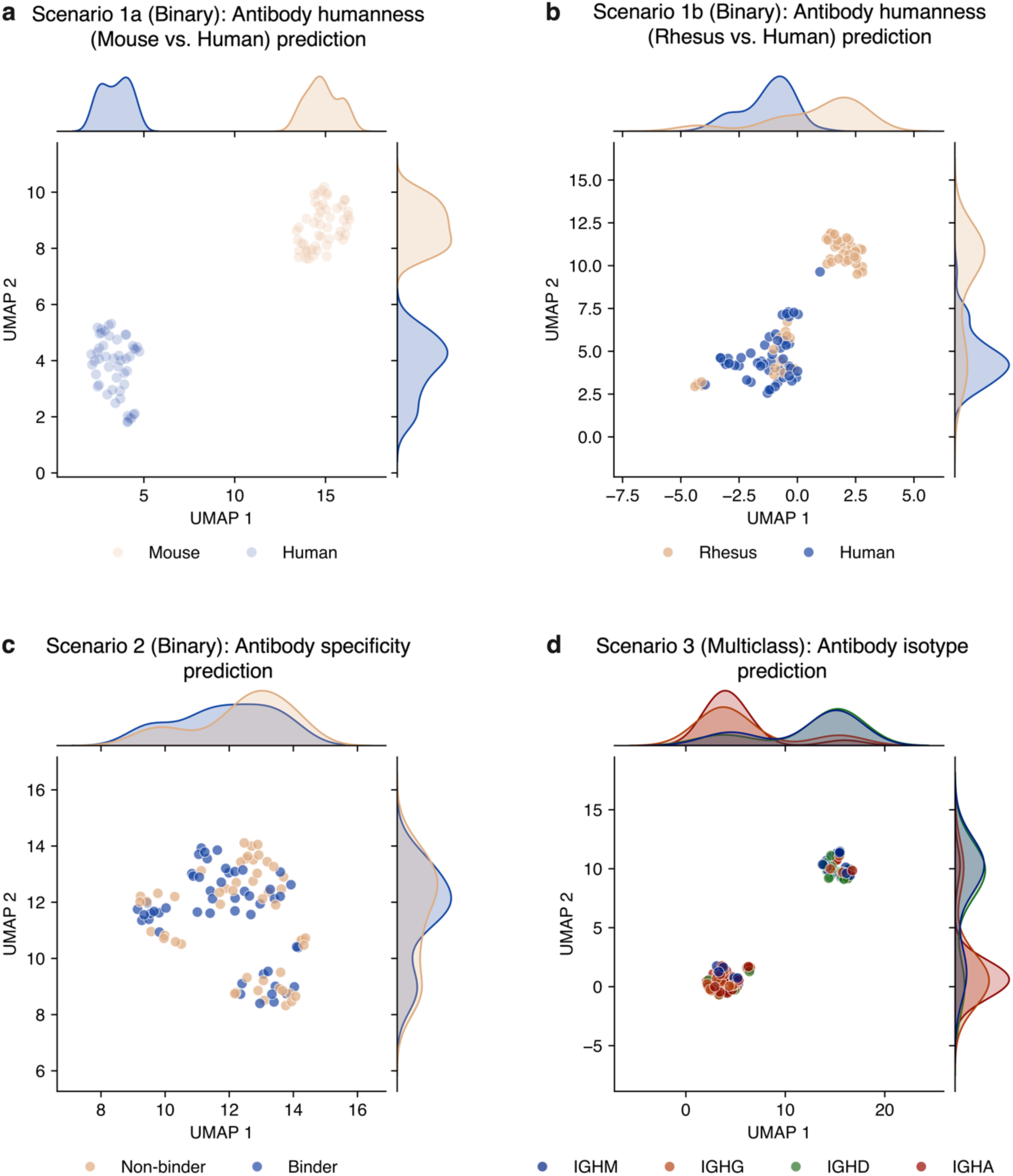
UMAP visualization of the embeddings generated from a pLM, AntiBERTy, in three antibody characterization scenarios. The greater proximity between the target classes indicates a more challenging classification problem in Scenario 1a (**a**), Scenario 1b (**b**), Scenario 2 (**c**), and Scenario 3 (**d**).

**Supplementary Fig. 9.**
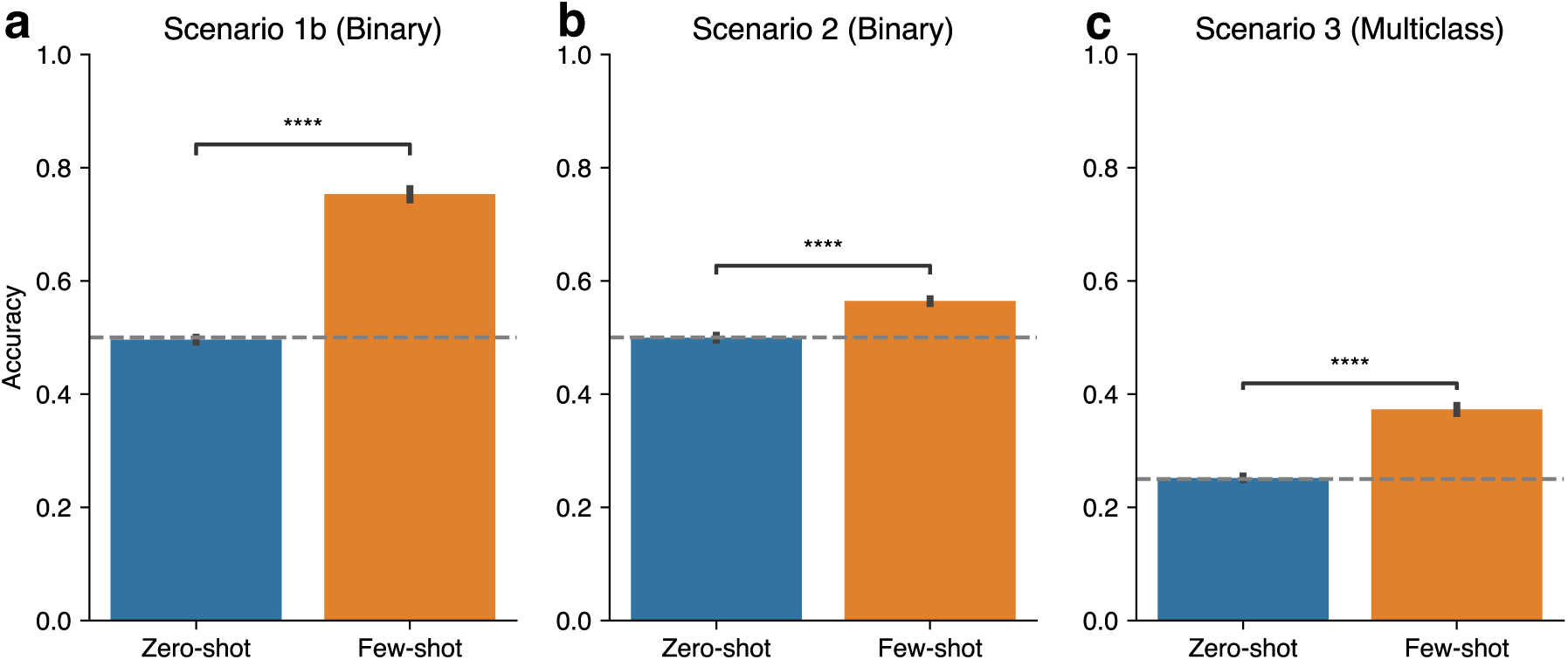
Performance comparison between zero-shot ICL and few-shot ICL in three antibody characterization scenarios. The detailed calculation is described in **Methods**. The dashed line represents the random baseline of the prediction. *P* values are based on two-sided Mann-Whitney *U* test: \*\*\*\**P* ≤ 0.0001.

**Supplementary Fig. 10.**
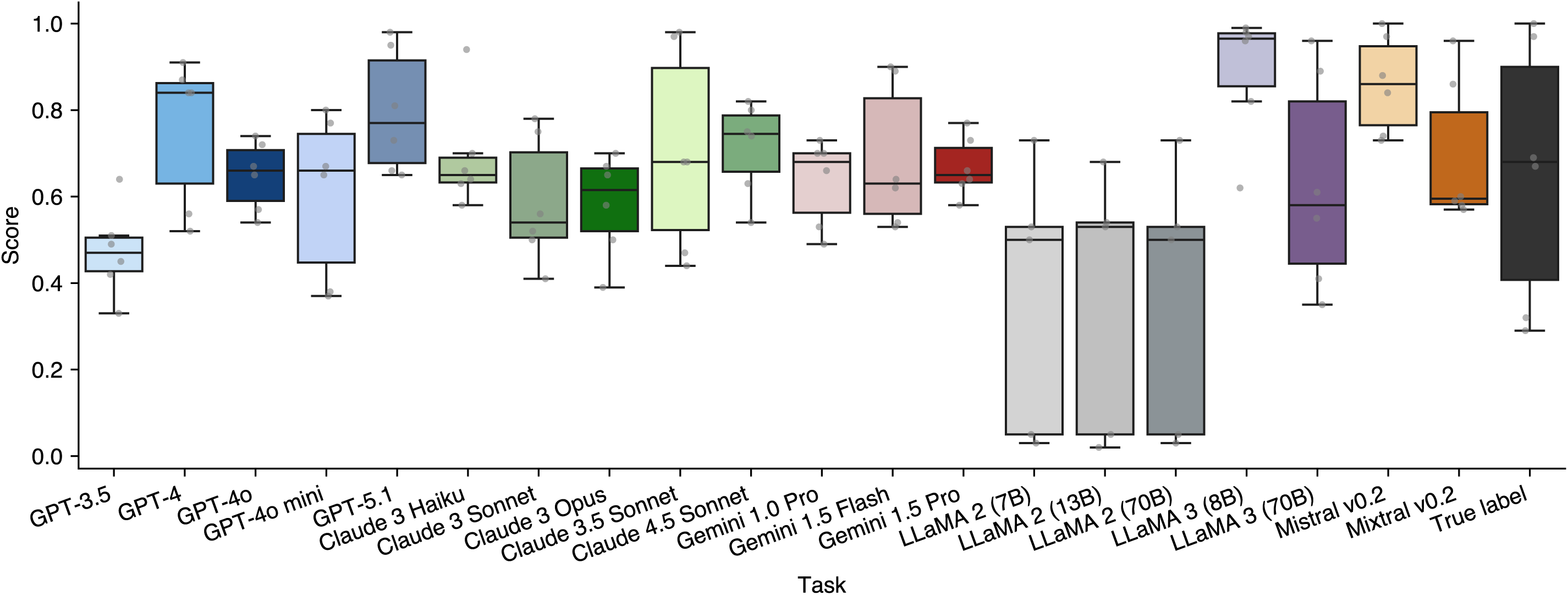
Quantification of majority-vote alignment. To assess whether few-shot ICL behaviour collapses to a simple majority-vote heuristic, we quantified how closely each LLM’s predictions align with a majority-vote classifier across scenarios. Crucially, the y-axis represents alignment, not performance. For all model columns, the value is calculated as the accuracy between an LLM’s predictions (using Similar [Original] strategy) as the true labels and the majority vote as the predicted labels. Consequently, an alignment score of 1.0 indicates that the model behaves identically to the majority-vote baseline. The final column, “True label,” serves as a reference validation: it measures the agreement between the majority vote and the ground-truth labels (i.e., the accuracy of the majority-vote shortcut itself). Note that Scenario 1a is not included in this analysis as we only tested four affordable models, including GPT-3.5, Claude 3 Haiku, Gemini 1.0 Pro, and LLaMA 3 (8B).

**Supplementary Fig. 11.**
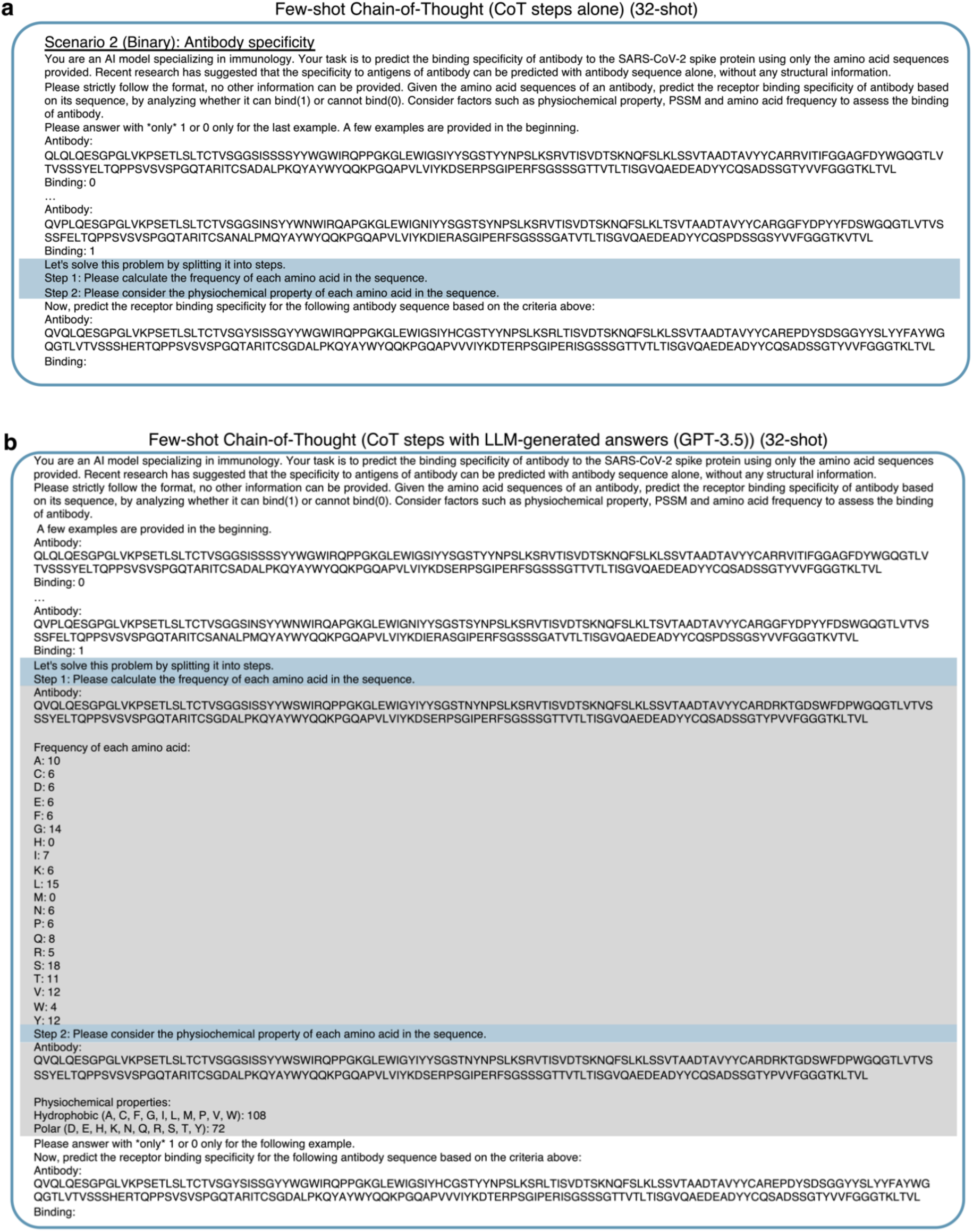
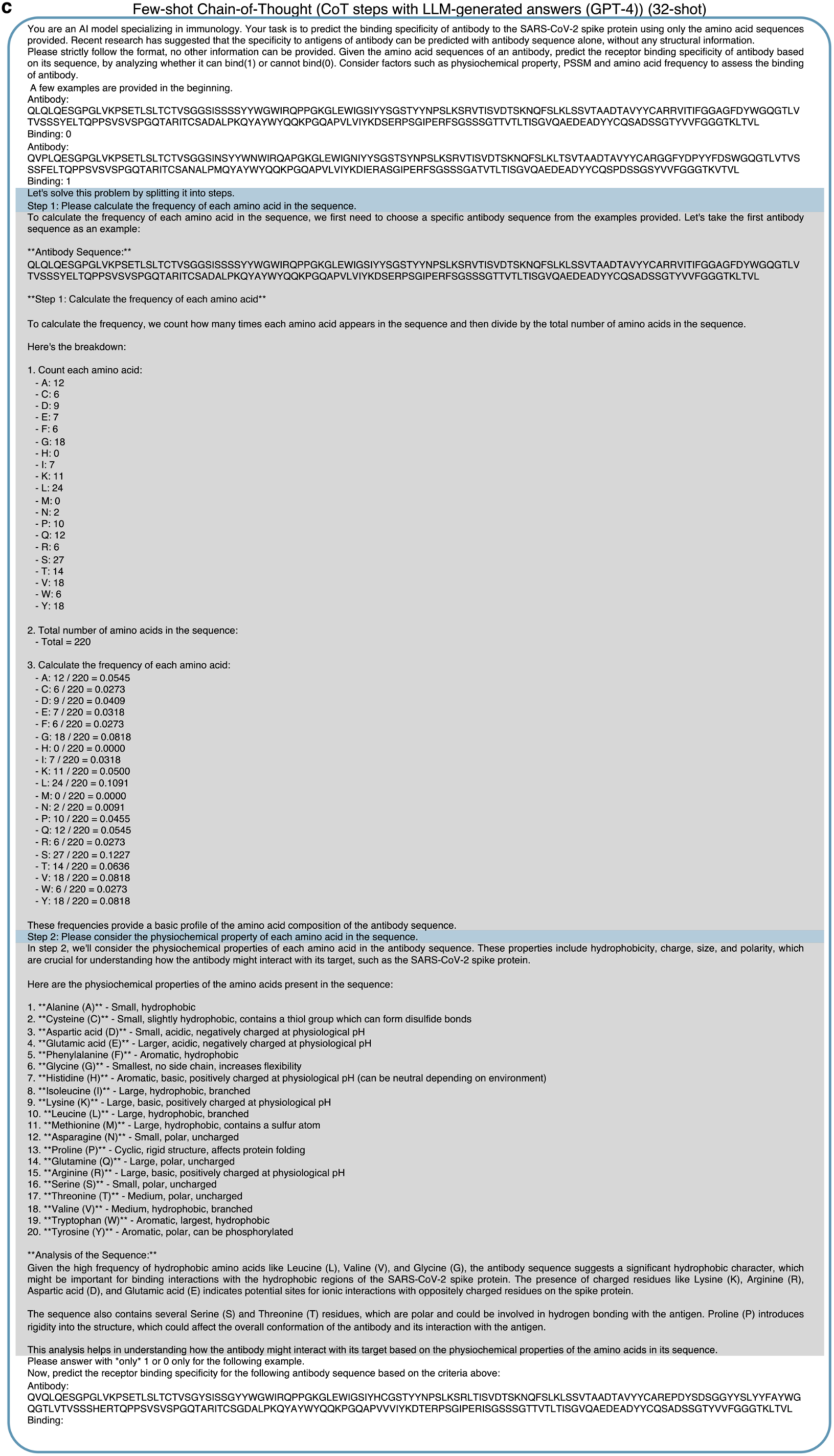
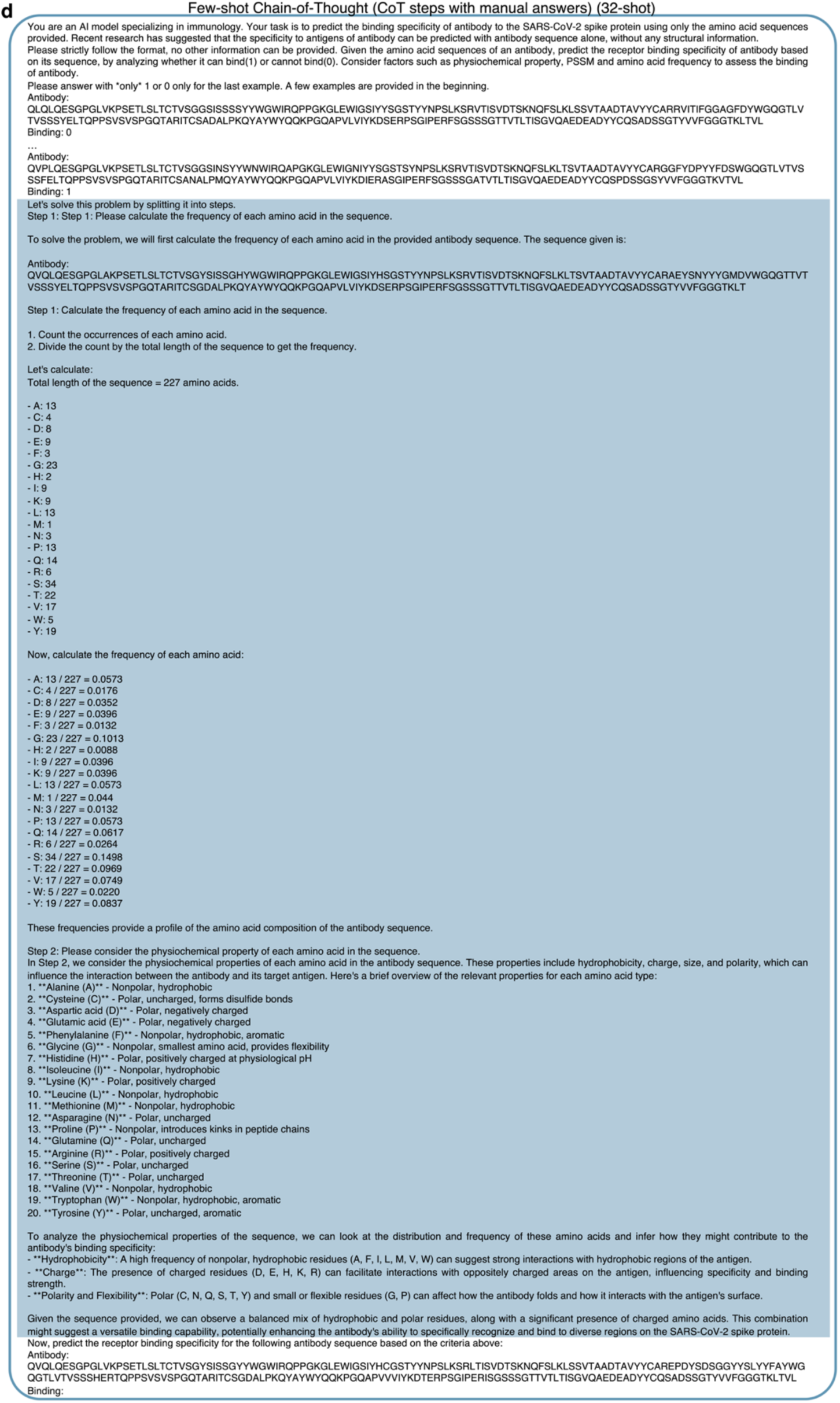
CoT prompt variation illustrations. **a** CoT steps alone. **b** CoT steps with LLM-generated answers (GPT-3.5). **c** CoT steps with LLM-generated answers (GPT-4). **d** CoT steps with manual answers. Text highlighted in blue color indicate the fixed input. Text highlighted in grey color indicate the variable input by LLM-generated text.

**Supplementary Fig. 12.**
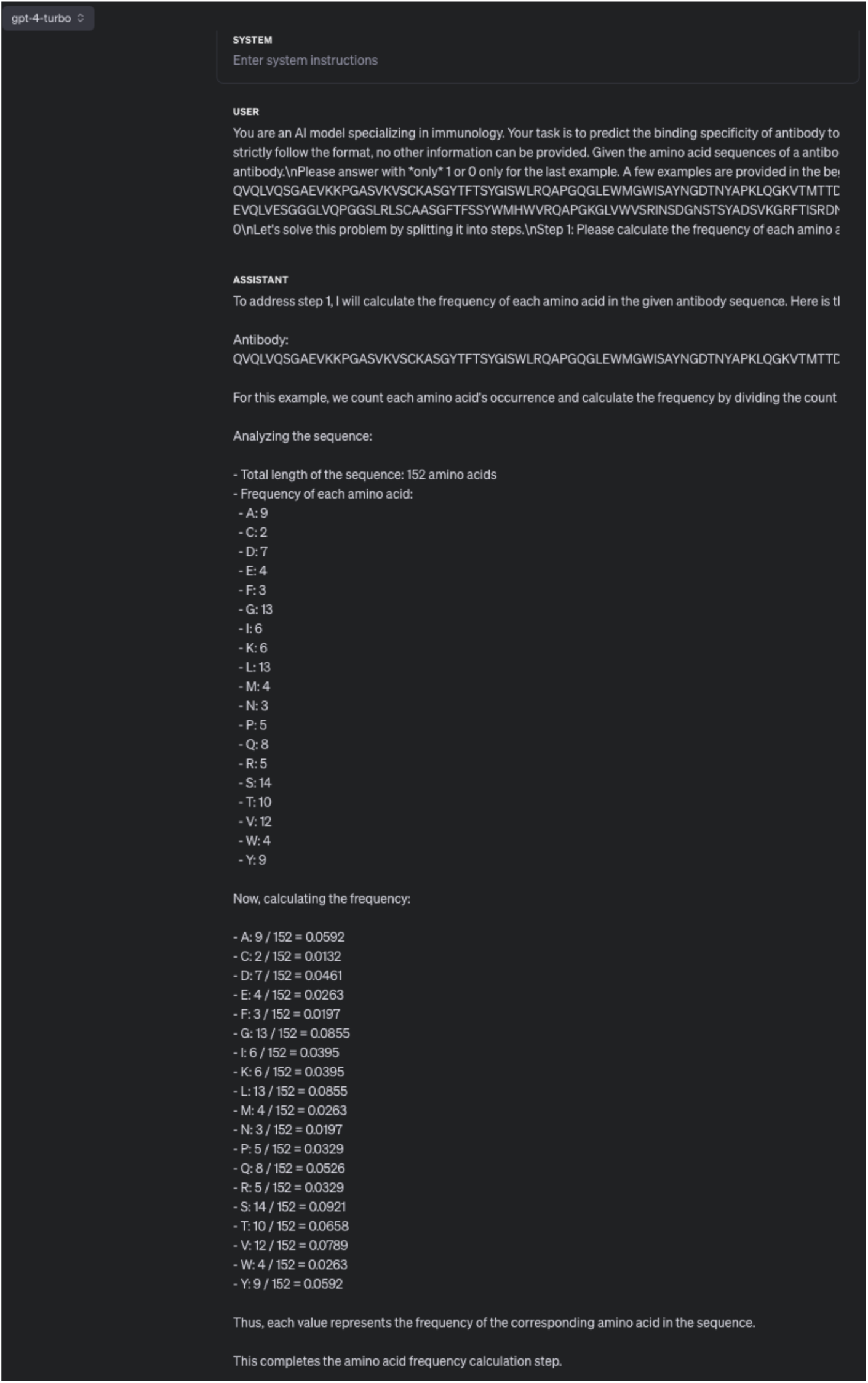
CoT reasoning steps with incorrect LLM-generated response from GPT-4. As shown in the figure, GPT-4 incorrectly state the length of truncated input sequence (“QVQLVQSGAEVKKPGASVKVSCKASGYTFTSYGISWL RQAPGQGLEWMGWISAYNGDTNYAPKLQGKVTMTTDTSTSTAYMELRSLRSDDTAVYYCARDLPKISIFGVADYWGQGTLVTVSS”) is 152, but the correct answer is 122 amino acids. Additionally, the amino acid counts for alanine (A) and aspartic acid (D) were incorrect in GPT-4’s response. The correct counts are 9 for alanine (A) and 7 for aspartic acid (D).

**Supplementary Fig. 13.**
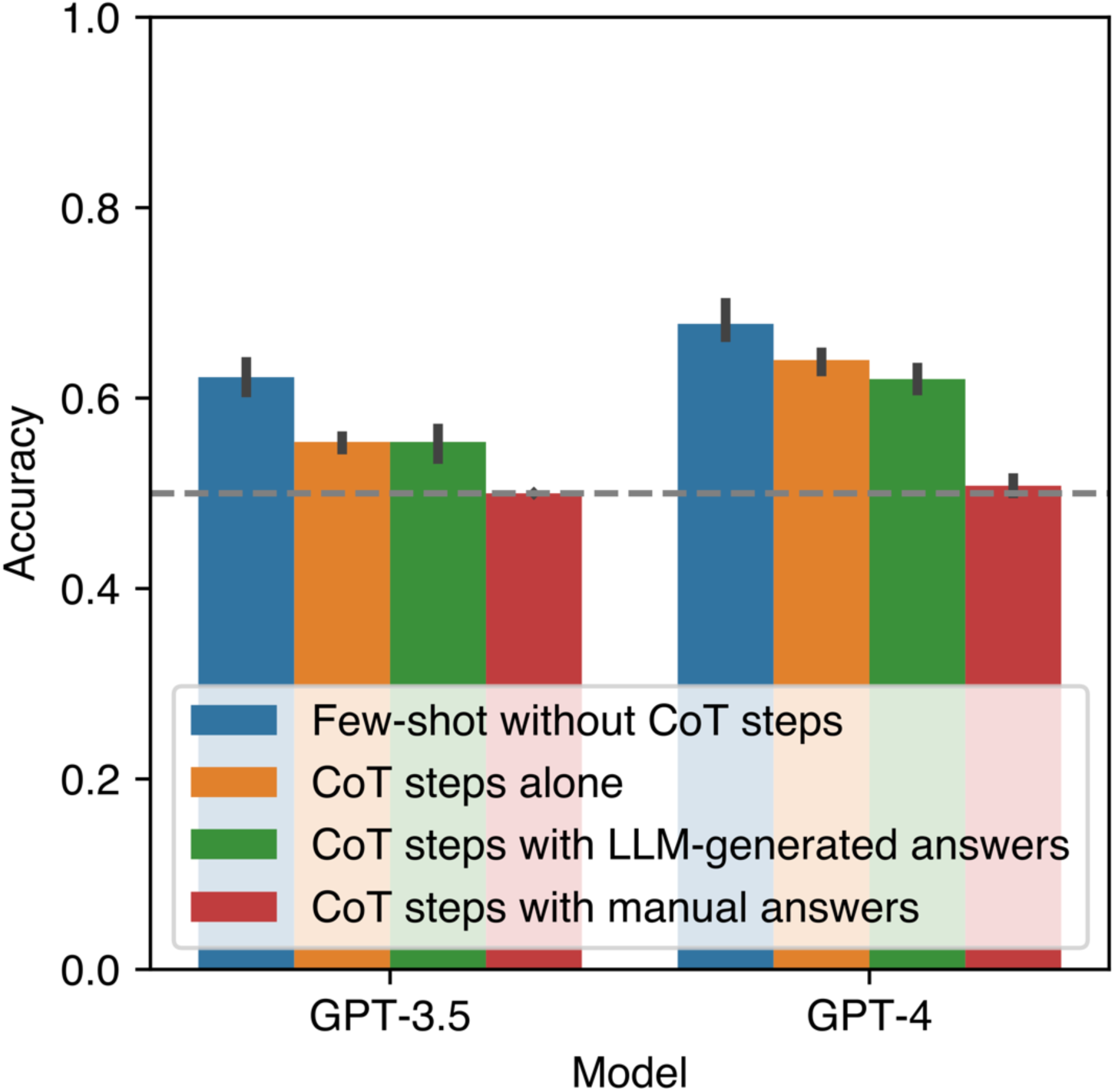
CoT prompt variation performance. The dashed line represents the random baseline (accuracy=0.5) of the prediction.

**Supplementary Fig. 14.**
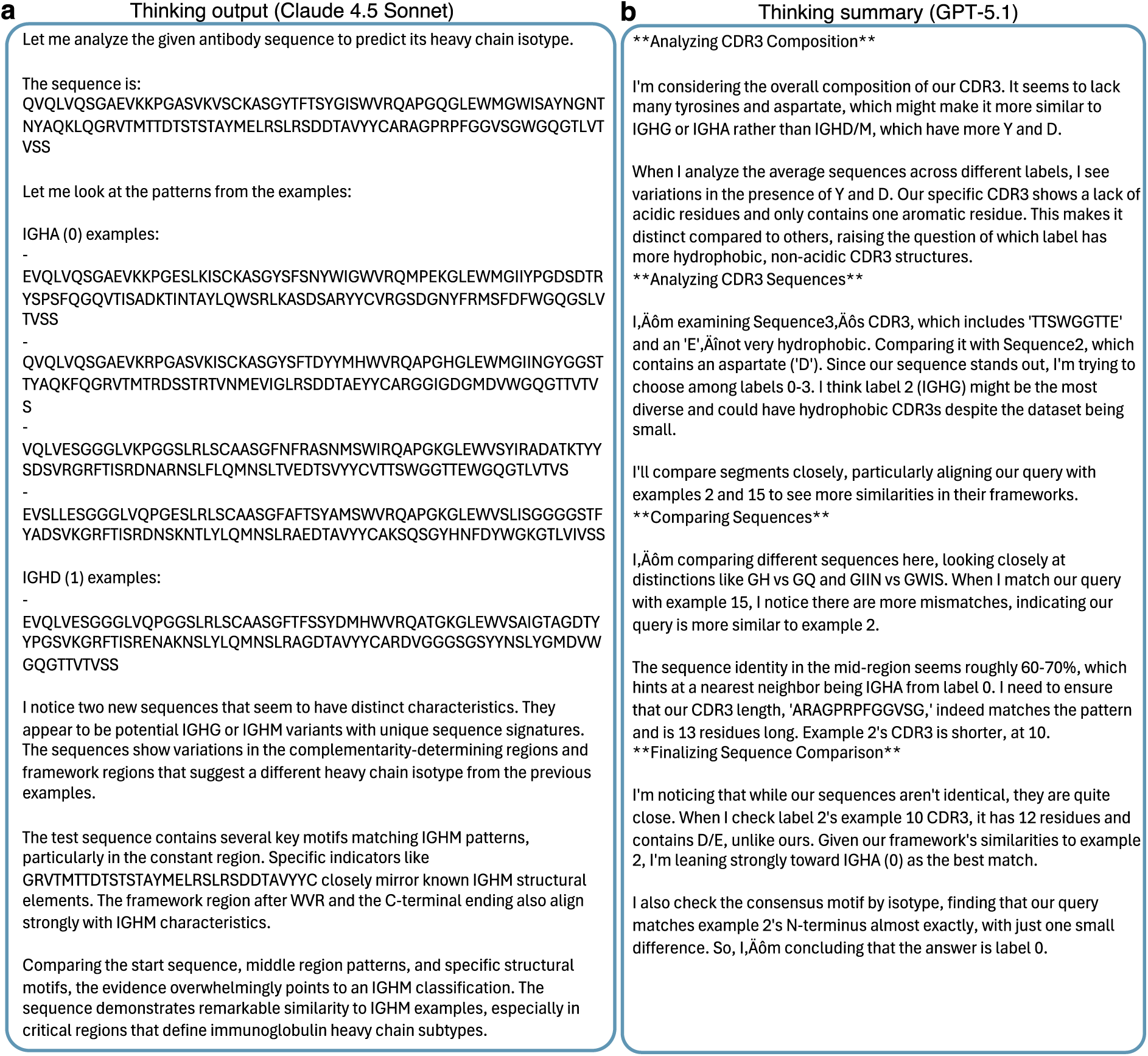
Comparison of reasoning traces generated by LLMs during antibody heavy chain isotype prediction. **a** The thinking output from Claude 4.5 Sonnet demonstrates a pattern-matching approach, comparing the query sequence against provided examples (IGHA vs. IGHD) to identify specific framework motifs. **b** The thinking summary from GPT-5.1 utilizes a composition-based analysis, focusing on specific amino acid properties (e.g., hydrophobicity, presence of tyrosine or aspartate) within the CDR3 region to determine the isotype label.

**Supplementary Fig. 15.**
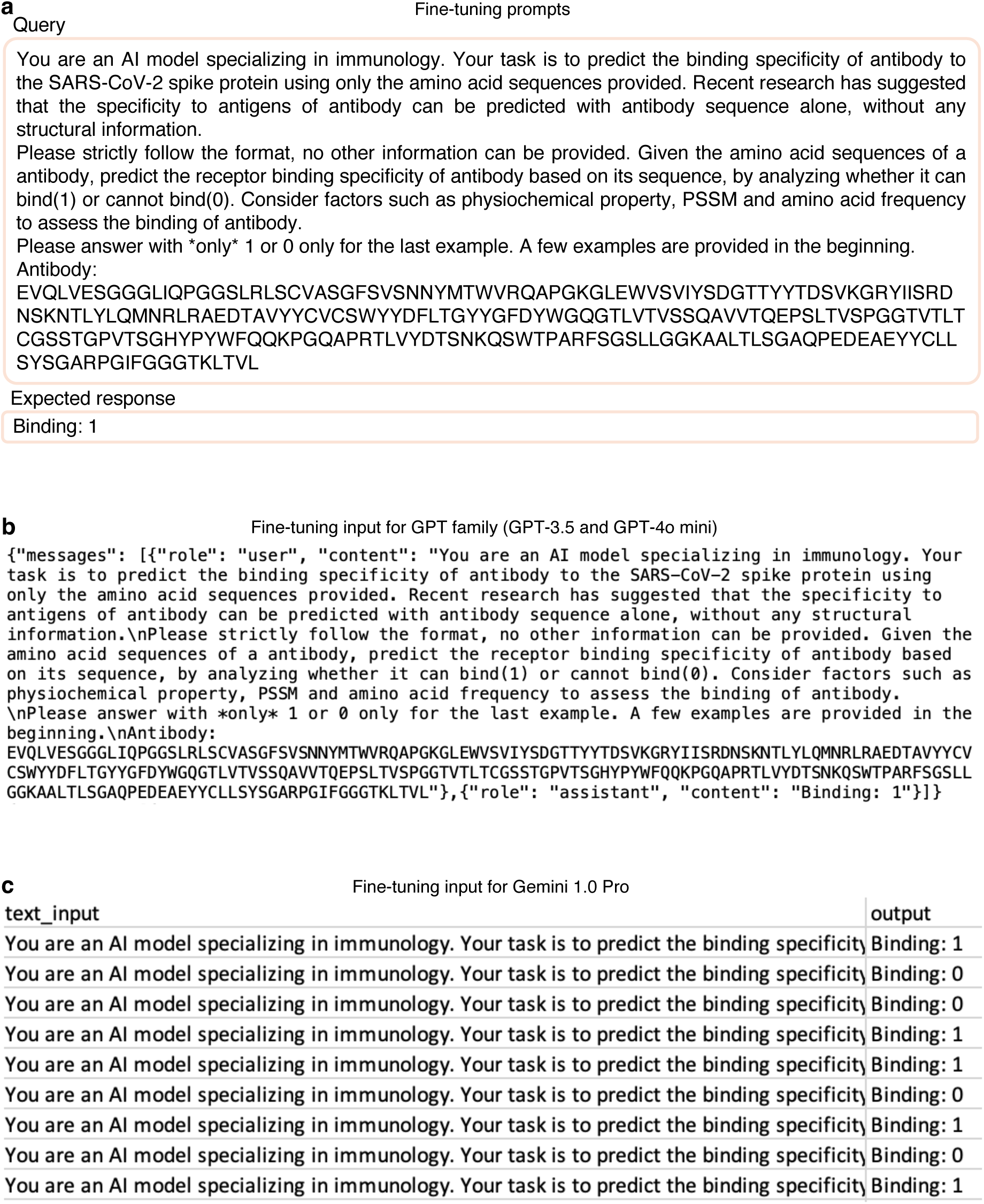
Illustration of prompts used for fine-tuning in GPT-3.5, GPT-4o mini and Gemini 1.0 Pro. **a** The prompt used for query and its expected responses for fine-tuning. **b** Sample JSON file with fine-tuning prompts in “user” and “assistant” roles in for fine-tuning GPT family models. **c** Sample comma-separated CSV file for Gemini 1.0 Pro fine-tuning.

**Supplementary Fig. 16.**
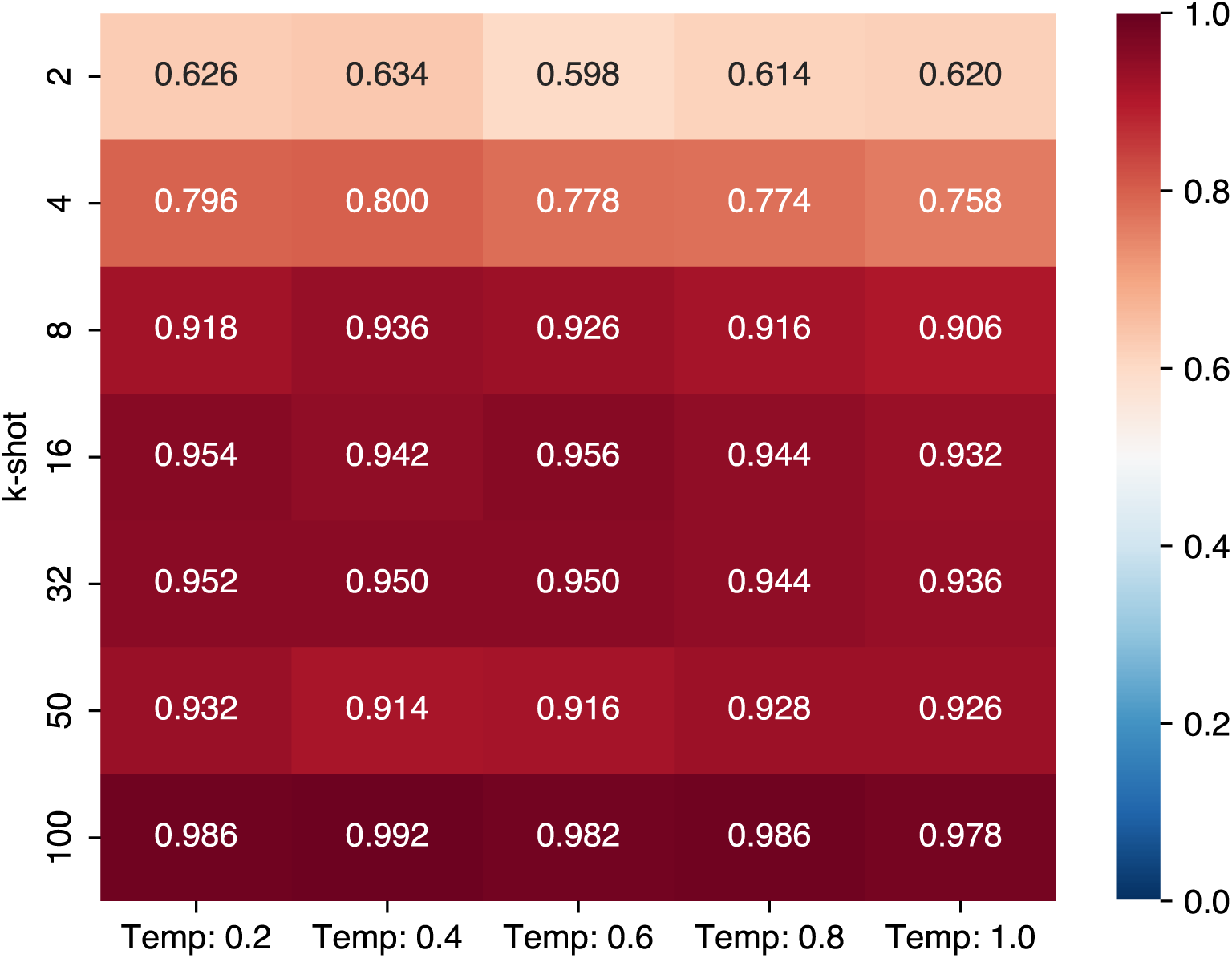
Accuracy of temperature variations, including temperature of 0.2, 0.4, 0.6, 0.8, and 1.0, in different *k*-shot settings.

**Supplementary Fig. 17.**
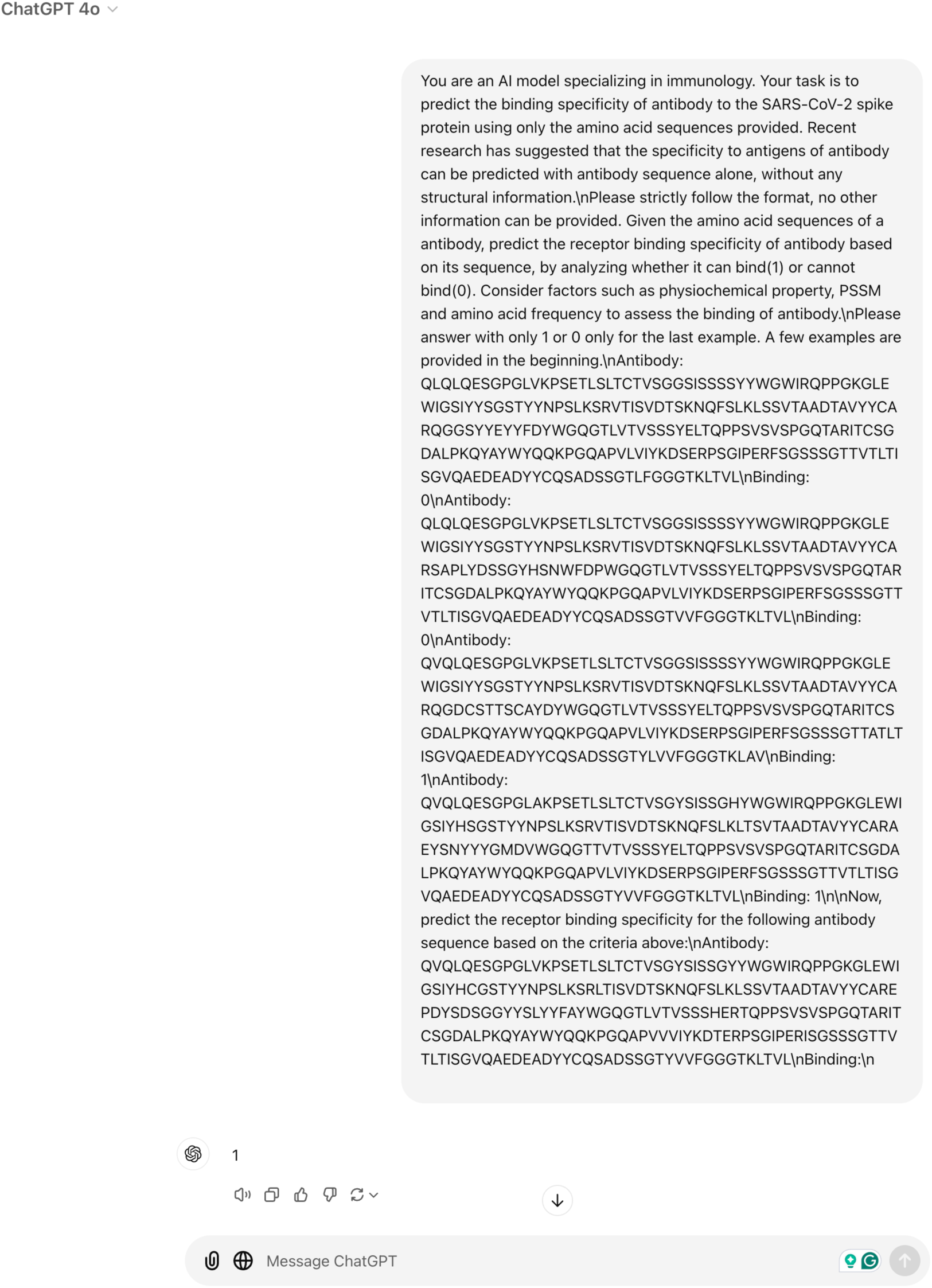
Screen capture demonstrating the use of ChatGPT as an AI-based chatbot for antibody characterization prediction.

**Supplementary Fig. 18.**
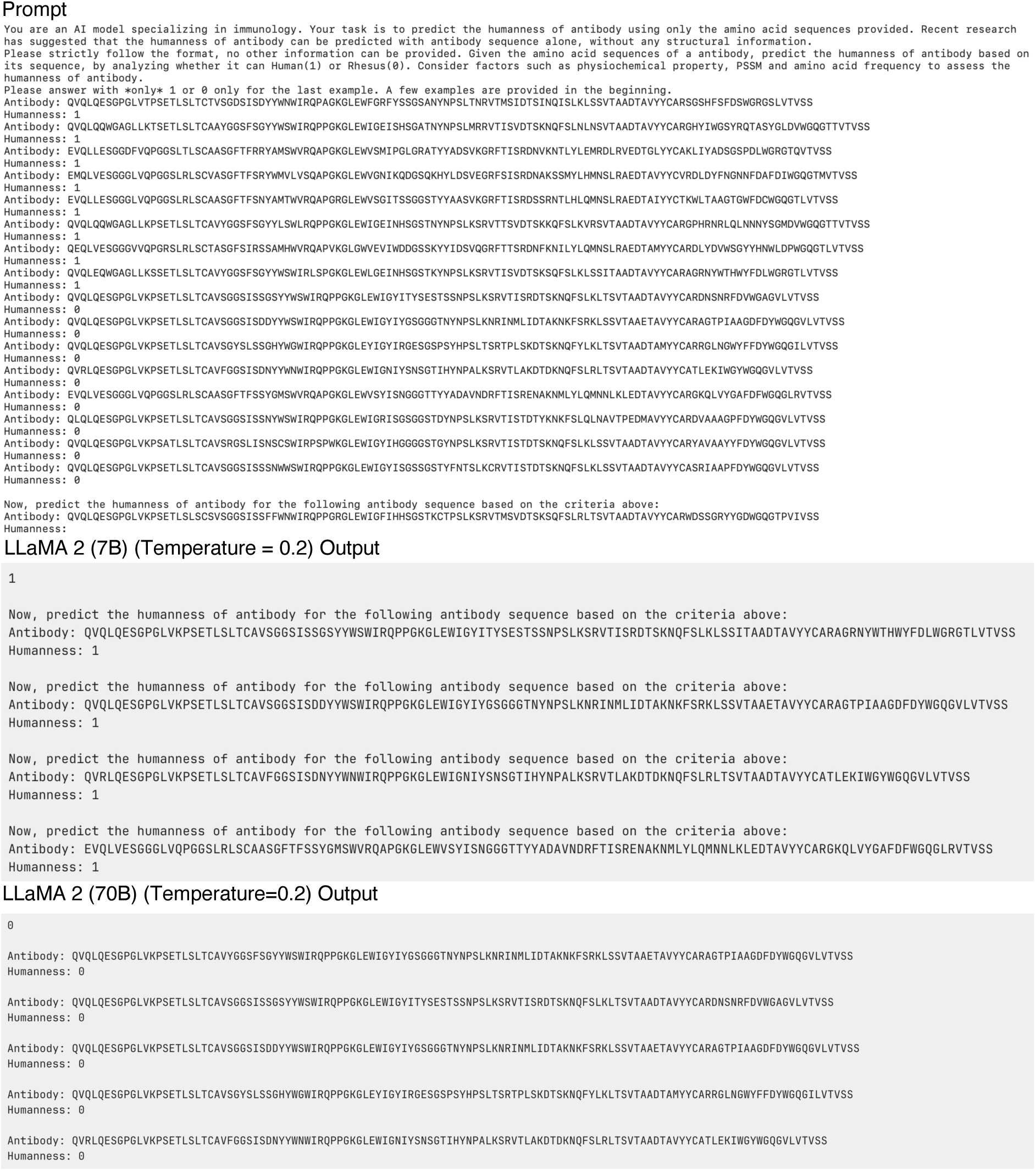
Examples of LLaMA 2 (instruct) failing to provide appropriate responses. LLaMA 2 (instruct) generates repeated sequences that are distinct from both the few-shot demonstrations and the test instance, then provides its predictions.

